# A metabolic model based on a pangenome core unveils new biochemical features of the phytopathogen *Xylella fastidiosa*

**DOI:** 10.64898/2026.03.23.713690

**Authors:** Paola Corbín-Agustí, Miguel Álvarez-Herrera, Miguel Román-Écija, Patricia Álvarez, Marta Tortajada, Blanca B. Landa, Juli Peretó

## Abstract

*Xylella fastidiosa* is a xylem-limited phytopathogenic bacterium responsible for severe diseases in many economically important crops. Despite its impact, its metabolism remains poorly characterized due to fastidious growth and the limited availability of defined culture media. Here, we reconstruct the first pangenome-based genome-scale metabolic model for *X. fastidiosa*, integrating conserved metabolic functions from 18 strains across five subspecies. The resulting core model, iXfcore, is manually curated and used to explore the species’ metabolic capabilities. Model simulations predict minimal nutritional requirements that guided us in the formulation of defined media supporting biofilm formation in vitro, providing validation of the model’s predictive capacity. Network analysis also identifies a previously undescribed pathway enabling growth on acetate as a sole carbon source. In addition, the model predicts the overproduction of polyamines, compounds linked to virulence in other phytopathogens. Experimental analyses confirm the production and secretion of polyamines in multiple *X. fastidiosa* strains, providing the first evidence of this capability. These results suggest that polyamine biosynthesis may represent an uncharacterized virulence factor for *X. fastidiosa*, potentially contributing to protection against host-induced oxidative stress. Overall, iXfcore provides a systems-level framework to investigate *X. fastidiosa* metabolism, generate testable hypotheses on its physiology and virulence, and support future strain-specific models and studies of host-pathogen metabolic interactions.

## 1. Introduction

*Xylella fastidiosa* is a fastidious, Gram-negative, xylem-limited phytopathogenic bacterium that infects more than 700 plant species worldwide (EFSA et al., 2026), including several crops of major agro-economic relevance. In some of these hosts, the bacterium causes devastating diseases such as Pierce’s disease in grapevine, citrus variegated chlorosis, almond leaf scorch, and olive quick decline syndrome (EPPO, 2023). The bacterium colonizes the xylem vessels of its plant hosts, and symptoms development depends on several bacterial virulence factors, particularly the formation of biofilm and host defense responses (Landa et al., 2022; De La Fuente et al., 2024). Biofilms also develop in the anterior foregut of sap-feeding insect vectors, which mediate bacterial transmission (Chatterjee et al., 2008).

*X. fastidiosa* is also characterized by substantial intraspecific diversity. Recent computational analyses support a revised taxonomy that groups the species into three major clades: clade I (corresponding to subsp. *pauca*), clade II (subsp. *multiplex*) and clade III (including three subsp.: *fastidiosa*, *morus* and *sandyi*) (Denancé et al., 2019). In parallel, multilocus sequence typing (MLST) has been widely used for the epidemiological characterization of strains (Scally et al., 2005) with 90 sequence types (STs) have been described to date (Jolley et al., 2018; EPPO 2023). Although STs do not provide genome-wide resolution and therefore cannot capture the full extent of gene-content variation, they remain useful for rapid strain typing, subspecies assignment, and initial epidemiological interpretation. However, the high recombination rates and the existence of some non-monophyletic STs raise concerns about the phylogenetic resolution and the robustness of current classification schemes based solely on MLST (Hanage et al., 2005; Vanhove et al., 2019; Landa et al., 2020). This genomic complexity suggests that marker-based classification alone is insufficient to infer functional traits, including metabolic capabilities that may influence host association, niche adaptation, and pathogenicity (Landa et al., 2022).

Despite decades of research, key aspects of *X. fastidiosa* physiology –particularly those linked to virulence, host adaptation, and nutritional requirements– remain poorly understood. This knowledge gap is especially relevant because the bacterium inhabits the xylem, a nutritionally restricted environment in which successful colonization, multiplication, and biofilm formation are expected to depend strongly on metabolic adaptation. Its inability to grow efficiently on standard bacteriological media, reflected in the species epithet “*fastidiosa*”, continues to be a defining feature of the species (Davis et al., 1981; Wells et al., 1987; Chatterjee et al., 2008). Empirical efforts have produced improved culture media that support faster growth (Davis et al., 1980; Wells et al., 1981; Davis et al., 1981; de Macedo Lemos et al., 2003), but none fully overcome its fastidiousness (Campanharo et al., 2003). Moreover, growth in currently available defined media remains slow and biomass production is limited. As a result, the metabolic basis of nutrient use, physiological specialization, and virulence-associated traits in *X. fastidiosa* is still only partially resolved (De La Fuente et al., 2024).

Genome-scale metabolic models (GEMs) are structured knowledge bases that integrate biochemical, genetic and genomic data into a system-level representation of the metabolic capabilities of an organism (O’Brien et al., 2015). Over the past two decades, GEMs have been applied in systems biology, enabling in silico prediction of genotype-phenotype relationships, reverse ecology paradigm to infer ecological features from genomic data, such as nutrient requirements prediction and growth medium design, and the discovery of metabolic network properties or pan-reactome comparisons (Yus et al., 2009; Monk et al., 2014; Gu et al., 2019; Ye et al., 2022; Qian & Ye, 2024). Constraint-based reconstruction and analysis (COBRA) is one of the foundational and most widely used approaches for working with GEMs. It achieves high predictive power by imposing physicochemical and environmental constraints (such as stoichiometry, thermodynamics and nutrient availability) that define a solution space encompassing all feasible metabolic states of the cell under different conditions (Ibarra et al., 2002; Palsson, 2015).

Here, we present an in silico approach to investigate key aspects of *X. fastidiosa* physiology through the reconstruction of a GEM. Although two metabolic models have previously been reconstructed for this phytopathogen, both are strain-specific (Gerlin et al., 2020; Oliveira et al., 2023). In this study, to account for its remarkable intraspecific genetic diversity while focusing on its conserved metabolic functions, a pangenome was assembled and used it as the template for a core metabolic model reconstruction. The resulting model–iXfcore–represents a species-level GEM that reflects metabolic functions conserved across *X. fastidiosa,* rather than traits restricted to particular subspecies or strains. Then, this reconstruction was experimentally validated by growth-phenotype profiling of a selection of *X. fastidiosa* strains. We also used the iXfcore model to design a defined minimal growth medium based on predicted nutritional requirements–an aspect that has received limited attention despite *X. fastidiosa* being the first phytopathogenic bacterium with a fully sequenced genome (Simpson et al., 2000). Beyond this, the metabolic network highlighted two notable features of the bacterial physiology. First, given the absence in *X. fastidiosa* of previously described acetate assimilation pathways, we sought a metabolic solution that would explain growth on acetate as the sole carbon source. That solution emerges from the combination of other metabolic submodules, providing genomic support for acetate utilization suggested by model-based analyses. Second, the iXfcore model predicts a remarkable capacity for polyamine biosynthesis. Polyamines are metabolites that are not only produced by plants as part of their oxidative stress and defense responses, but have also been previously linked to virulence in other phytopathogens such as *Ralstonia solanacearum* (Lowe-Power et al., 2018) and *Pseudomonas syringae* (Solmi et al., 2025). In this study, we experimentally confirmed polyamine production in *X. fastidiosa* for the first time, suggesting a possible role in host-pathogen interactions. Together, this work delivers the first species-level GEM of *X. fastidiosa* and establishes a systems biology framework to investigate its physiology, nutritional requirements, and metabolic determinants of pathogenicity.

## 2. Materials and Methods

### 2.1. Genome data preparation and pangenome reconstruction

A total of 18 *X. fastidiosa* strains belonging to five subspecies were considered for the pangenome assembly (Table 1). Complete, chromosomal, whole-genome data was retrieved from the NCBI Genome database (Sayers et al., 2024). For each strain, the associated plasmid sequences, when available, were also inspected using the BlastKOALA annotation server (Kanehisa et al., 2016) and the NCBI annotation. As none of the plasmid-encoded genes were associated with relevant metabolic functions, plasmid sequences were excluded from the pangenome analysis. Coding sequences, excluding those annotated as pseudogenes, were clustered into ortholog groups (OGs) using InParanoid (Remm et al., 2001), applying a minimum of 80 % sequence identity and at least 50 % pairwise aligned sequence length, using the BLOSUM 80 substitution matrix. Pairwise OGs generated by InParanoid were merged into multi-strain OGs using QuickParanoid (http://pl.postech.ac.kr/QuickParanoid/). Genes without a best bidirectional hit in any other genome were classified as “unique” while “accessory” genes were defined as those present at least in two strains. Genes belonging to OGs shared across all strains were considered “core”. Core, accessory and unique gene sets were annotated with BlastKOALA automatic annotated server to explore the functional traits.

**Table 1.** *Xylella fastidiosa s*trains used for the pangenome reconstruction and genome-scale metabolic modeling. The complete genomes of eighteen strains of *X. fastidiosa* belonging to five subspecies of the phytopathogen were used for the reconstruction of the pangenome. For each case, the strain identifier, the subspecies to which it belongs, the host, the site of isolation, the disease it causes and the NCBI accession number for the genome are indicated.

| Identifier | Subspecies | Host | Location | Disease<sup>a</sup> | ID RefSeq |
| --- | --- | --- | --- | --- | --- |
| Temecula1 | *fastidiosa* | *Vitis vinifera* | California, USA | PD | NC_004556.1 |
| M23 | *fastidiosa* | *Prunus dulcis* | California, USA | PD, ALS | NC_010577.1 |
| MUL0034 | *morus* | *Morus alba* | California, USA | MLS | NZ_CP006740.1 |
| M12 | *multiplex* | *Prunus dulcis* | California, USA | ALS | NC_010513.1 |
| ESVL | *multiplex* | *Prunus dulcis* | Alacant, Spain | ALS | NZ_CP099516.1 |
| IVIA5901 | *multiplex* | *Prunus dulcis* | Alacant, Spain | ALS | NZ_CP047134.1 |
| XYL468 | *multiplex* | *Olea europaea*<br>*var. sylvestris* | Mallorca, Spain | OQDS | NZ_CP136966.1 |
| CFBP8418 | *multiplex* | *Spartium junceum* | Corse, France | LS | NZ_CP136979.1 |
| 9a5c | *pauca* | *Citrus sinensis* | São Paulo, Brazil | CVC | NC_002488.3 |
| U24d | *pauca* | *Citrus sinensis* | São Paulo, Brazil | CVC | NZ_CP009790.1 |
| J1a12 | *pauca* | *Citrus sinensis* | São Paulo, Brazil | Non-pathogenic | NZ_CP009823.1 |
| Pr8x | *pauca* | *Prunus domestica* | São Paulo, Brazil | LS | NZ_CP009826.1 |
| 3124 | *pauca* | *Coffee arabica* | São Paulo, Brazil | CLS | NZ_CP009829.1 |
| Hib4 | *pauca* | *Hibiscus fragilis* | Distrito Federal, Brazil | LS | NZ_CP009885.1 |
| Salento-2 | *pauca* | *Olea europaea* | Puglia, Italy | OQDS | NZ_CP016610.1 |
| De Donno | *pauca* | *Olea europaea* | Puglia, Italy | OQDS | NZ_CP020870.1 |
| XYL1961/18 | *pauca* | *Olea europaea*<br>*var. europaea* | Eivissa, Spain | OQDS | CP171775.1 |
| Ann-1 | *sandyi* | *Nerium oleander* | California, USA | OLS | NZ_CP006696.1 |

<sup>a</sup> PD: Pierce's Disease; ALS: Almond Leaf Scorch; MLS: Mulberry Leaf Scorch; CVC: Citrus Variegated Chlorosis; CLS: Coffee Leaf Scorch; OQDS: Olive Quick Decline Syndrome; OLS: Oleander Leaf Scorch; LS: Leaf Scorch.

### 2.2. Metabolic model reconstruction and refinement

For COBRA, established protocols were followed (Thiele & Palsson, 2010; Nogales, 2017). An initial draft reconstruction of the metabolic network was generated based on the core gene set of the *X. fastidiosa* 9a5c strain using the automatic workflow provided by ModelSEED (Overbeek et al., 2005; Henry et al., 2010). In addition, draft reconstructions were generated for all remaining strains to support the manual curation of the core draft model. Curated genome-scale metabolic models of *Escherichia coli* iJO1366 (Orth et al., 2011) obtained from BiGG Models (King et al., 2016), *X. fastidiosa* Xfm1158 (Gerlin et al., 2020) and iMS508 (Oliveira et al., 2023) from BioModels (Malik-Sheriff et al., 2020), and *Xanthomonas campestris* pv. *campestris* str. B100 (Schatschneider et al., 2013) were used as references during the curation process. Subsequently, the model was iteratively refined to ensure consistency with available physiological and biochemical data (see Supplementary Methods: Metabolic model refinement).

In parallel with the rest of the network, the biomass reaction was also curated, using as template the one generated by ModelSEED. The suitability of each metabolite in the template and potential biomass components were evaluated, including the molar contribution of each one, based on biochemical data and the biomass reactions from the reference models. Manipulation of metabolic reconstructions was carried out with the COBRApy library in Python (Ebrahim et al., 2013). NetworkX v2.4 (Hagberg et al., 2008), Escher v1.7.3 (King et al., 2015), yED Graph Editor v3.18.2 (yWorks GmbH, 2020) and Cytoscape v3.7.1 (Shannon et al., 2003) were used for metabolic network visualization during the curation process. The quality of the final model was evaluated using the MEMOTE v.0.13.0 tool (Lieven et al., 2020). The final GEM is provided in JSON, MAT-File and Systems Biology Markup Language (SBML) formats (Supplementary Material S1).

### 2.3. In silico constraints and growth conditions

For reversible reactions, upper and lower bounds of 1000 and -1000 mmol·gDW^-1^·h^-1^, were respectively applied. For irreversible reactions, the lower (or upper) bound was fixed at 0 mmol·gDW^-1^·h^-1^. Maximization of growth rate was set as the objective function, leaving oxygen exchange unconstrained. The growth-associated maintenance energy (GAM) was fixed at 50.12 mmol·gDW⁻¹·h⁻¹, while the non-growth-associated maintenance energy (NGAM) was set to 3.073 mmol·gDW⁻¹·h⁻¹, according to Gerlin et al. (2020).

For simulation of growth conditions, the lower bounds of exchange reactions were set to zero except for the inorganic ions included in the predicted minimal medium. For minimal medium simulations, glutamine uptake flux was constrained to -7.25 mmol·gDW^-1^·h^-1^ as the carbon and nitrogen source, based on environmental, biochemical and previous metabolic modeling data (Davis, 1992; Gerlin et al., 2020; Orth et al., 2011). For trade-off simulations, glutamine uptake was fixed at -15.665 mmol·gDW^-1^·h^-1^. More information can be found in Supplementary Methods: In silico constraints and growth conditions.

### 2.4. Flux Balance Analyses

Flux Balance Analysis (FBA) and parsimonious Flux Balance Analysis (pFBA) were performed using the COBRApy software (Ebrahim et al., 2013). The optimization problems were solved with the GNU Linear Programming Kit (GLPK) v.5.0 or the IBM ILOG CPLEX Optimizer v.22.1.1. FBA and pFBA were used to compute the maximum value of the objective function and the corresponding flux distribution (Orth et al., 2010; Thiele & Palsson, 2010).

### 2.5. Design of synthetic defined minimal media

To design a model-driven minimal medium sustaining the FBA-predicted growth rate under the fixed L-glutamine uptake constraint, the number of active exchange reactions was minimized. This was formulated as a mixed-integer linear programming (MILP) problem and solved using the GLPK solver, employing the *minimal_medium* implementation from the COBRApy medium module (Ebrahim et al., 2013). Based on the predicted metabolite requirements, six synthetic defined minimal media were designed (mXC1-mXC6, Table S1). The initial formulations (mXC1 and mXC2) were derived from the in silico minimal medium, using glutamine as carbon and nitrogen source, and different iron sources. For subsequent formulations (mXC3 to mXC6), glutamine was replaced with equivalent quantities of alternative carbon sources (sodium acetate or ribose), supplementing with nitrogen sources. Generally, ammonium sulfate was used as an inorganic nitrogen source, whereas cysteine was used as an alternative nitrogen source in mXC6 formulation. To validate the biotin-related predictions under growth on acetate, mXC4 medium was supplemented with biotin. Concentrations of all components were adapted from existing media known to support *X. fastidiosa* growth. Concentrations of glutamine, hemin chloride and inorganic salts were adapted from PW synthetic medium (Davis et al., 1981), while ferric pyrophosphate concentrations were derived from BCYE (Wells et al., 1981) and 3G10R media (Leite et al., 2004). Concentration of (NH_4_)_2_SO_4_ was obtained from XVM2 medium (Hiery et al., 2013), cysteine from BCYE and XF-26 (Chang, 1993), and biotin from SXM (Neumann, 2003). All media were adjusted to a pH of 6.9 and brought to a final volume of 1 L with sterile distilled water. PD3 broth (Davis et al., 1981) was selected as the reference medium because it is widely used for routine cultivation of *X. fastidiosa*, and served as a positive growth control.

### 2.6. Acetate assimilation pathways research

To investigate the metabolic strategy employed by *X. fastidiosa* for the conversion of acetate into biomass, the diverse metabolic assimilation pathways described in the literature were searched against the *X. fastidiosa* genome. Reference protein sequences corresponding to the enzymatic steps involved in each pathway were retrieved from the BRENDA database based on their associated EC numbers. Reciprocal BLASTP (v2.9.0+) (Camacho et al., 2009) searches were conducted between these reference sequences and the annotated amino acid sequences of *X. fastidiosa*. Substantial outcomes were obtained by setting specific filtering parameters (coverage greater than 80 %, e-value ≤ 10^−6^). Additionally, hmmscan, hmmsearch and pfamscan (Madeira et al., 2024) were used to identify conserved protein domains and domain architectures, providing additional support for functional inference. Finally, the results were manually revised and some selected candidates and enzymatic activities were reviewed using the UniProt database (UniProt Consortium, 2025) to validate results.

### 2.7. Polyamine and virulence factors trade-off prediction

Polyamine biosynthesis and degradation pathways were manually curated and revised as described in the Supplementary Methods metabolic model refinement section. Special attention was paid to S-adenosylmethionine (SAM) regeneration and the L-methionine salvage pathway. As no experimentally validated polyamine export mechanism has been described in *X. fastidiosa*, demand reactions for putrescine, spermidine, and spermine were incorporated to simulate intracellular production and secretion potential.

To investigate potential trade-offs between biomass production, polyamine secretion and virulence factors synthesis (EPS and LesA protein), a two-step optimization strategy was applied. First, an experimental-like scenario was reproduced constraining biomass production (0.1608 h⁻¹), EPS biosynthesis (1.296 mmol·gDW⁻¹·h⁻¹) and LesA production (0.0024 mmol·gDW⁻¹·h⁻¹) as described in Gerlin et al. (2020). Under these constraints, glutamine uptake was minimized, resulting in a flux of -15.665 mmol·gDW⁻¹·h⁻¹. This value was used as the carbon and nitrogen source uptake for all trade-off simulations. In the second step, biomass was constrained to a minimum of 0.1608 h⁻¹, and the feasible production space of the three target metabolites was explored. For each metabolic product, the maximum achievable flux at the minimum fixed biomass level was determined by FBA by maximizing the corresponding demand or export reaction. Uniform grids (30 intervals per axis) ranging from zero to the maximum achievable flux were generated for each metabolite. For every combination of target fluxes, biomass was constrained to a minimum value of 0.1608 h⁻¹ and the model was re-optimized maximizing the growth rate as the objective function using the CPLEX Optimizer. Only feasible solutions were retained. This sampling enabled the reconstruction of two-dimensional phase planes approximating the surface of a Pareto front, representing the trade-off between metabolite production and biomass growth rate. Phase planes were obtained by fixing one product to zero and projecting the feasible region of the remaining two metabolites. The resulting solution space was visualized using tricontour and tricontourf functions from the Matplotlib library. Biomass flux was represented as a continuous color gradient.

### 2.8. Bacterial strains and culture conditions

Seven *X. fastidiosa* strains were used in the different in vitro studies: two from subspecies *fastidiosa* isolated from grapevine in California (Temecula1) and Mallorca, Spain (IVIA5770); three from subspecies *multiplex* isolated from almond trees in Alacant, Spain (ESVL and IVIA5901) and wild olive in Mallorca (XYL468); and two from subspecies *pauca* isolated from olive trees (De Donno and XYL1961/18) in Puglia, Italy, and Eivissa, Spain, respectively. Stocks of *X. fastidiosa* strains were stored in phosphate buffered saline (PBS) plus 50 % glycerol at -80 °C. For all the experiments, strains were recovered from glycerol stocks on plates of PD3 (Davis et al., 1981) or BCYE (Wells et al., 1981) media incubated at 28 °C in the dark. After 6 to 8 days, cells were re-streaked onto new plates and cultured for 5 to 7 days before using them in each experiment.

### 2.9. Growth of *Xylella fastidiosa* strains in synthetic defined media

The potential utilization of 190 carbon sources of five *X. fastidiosa* strains (ESVL, De Donno, IVIA5901, Temecula1, and XYL1961/18) was determined using the PM1 and PM2 Biolog phenotype microarray (PM) system (Biolog, Hayward, California) (Supplementary Methods-Phenotypic characterization of *Xylella fastidiosa* strains).

In vitro culture experiments using the six minimal media described in section 2.5 (Table S1) were conducted with the seven *X. fastidiosa* strains mentioned in section 2.8 as previously described by Román-Écija and coworkers (2023), to assess the ability of the designed minimal media to support bacterial growth, as well as biofilm formation. Briefly, a cell suspension of each strain was diluted in each medium to reach at OD_600_ = 0.08. Cell suspensions (200 µL) were inoculated in polystyrene 96-well microplates (Sarstedt, Nümbrecht, Germany) and incubated at 28 °C during 7 days without agitation. Total bacterial growth was measured daily by measuring OD_600_. After the 7-days incubation, planktonic phase was discarded and biofilm formation was estimated using the crystal violet assay. At least four independent experiments were conducted, and each experiment included five replicated wells per treatment. Non-inoculated medium served as negative control.

### 2.10. Polyamines production by *Xylella fastidiosa* strains in liquid medium

To validate the metabolic model predictions for the ability of *X. fastidiosa* to produce polyamines during in vitro culture, two experiments were performed. An initial experiment was carried out to determine whether *X. fastidiosa* produces polyamines. For that, 10 mL of cell suspensions at OD_600_ = 0.05 of six *X. fastidiosa* strains (those mentioned in section 2.8, except for IVIA5901) were used to inoculate 50-mL glass flasks and incubated under agitation at 28 °C. After 9 days of growth, two types of samples were collected for polyamine extraction. The first one included *X. fastidiosa* cells from planktonic growth plus those present in the biofilm formed on the glass surface at the air-liquid interface. For that, cells were scratched from the attachment ring formed on the glass wall and vortexed. Then, the bacterial cell pellet (CP) was obtained after centrifugation at 14,000 rpm for 10 min of the entire culture. The second type of sample included the culture supernatant (SP). At least two biological replicates for each strain were performed. For the second experiment one strain per subspecies was selected, namely, ESVL, De Donno, and Temecula1. In this experiment, three types of samples were obtained after 5 and 12 days of growth. For each strain and sampling point, culture cells in planktonic growth were collected by transferring the entire culture to a new falcon tube. Then, a cell pellet from planktonic growth (CP-P) and a SP were obtained after centrifugation as described above. To collect cells growing in biofilms, the biofilm formed on the glass surface was scraped with a loop and resuspended with a pipette using 1 mL of sterile distilled water. Then, a cell pellet from biofilms (CP-B) was collected by centrifugation and removal of the supernatant. In this second experiment, four biological replicates for each strain and condition were obtained.

The polyamine extraction and quantification of CP-P, CP-B, and SP samples were carried out according to Solé-Gil and coworkers (2019) with small modifications. Briefly, bacterial cell pellets were resuspended in 1 mL sterile water, treated with lysozyme (10 mg·mL^-1^), and kept on ice for 10 min. Subsequently, bacterial cells were disrupted by sonication (10 cycles of 30 s each). After centrifugation at 10,000 rpm for 10 min, 500 µL of the supernatants were treated with 2.5 mL of perchloric acid 5 % (w/v). Culture supernatants (2.5 mL) were also treated with 2.5 mL of perchloric acid. At this point, pre-treated samples were stored at -20 °C and sent for polyamine analysis to the Metabolomics Service of the Institute of Molecular and Cellular Biology of Plants (IBMCP, Valencia). Before analysis, 500 µL of the internal standard (1,6-hexanediamine 1 mM) were added to each sample. Polyamines were quantified by HPLC analysis as described by Santos and coworkers (2021) in triplicate.

### 2.11. Data processing and statistical analysis

To evaluate growth in minimal media, growth curves were obtained by accumulating OD_600_ values over time from the date of inoculation. The OD_600_ value of the control wells was subtracted from each time point to obtain the net absorbance. Cumulative growth was then calculated by subtracting the initial net absorbance at the beginning of the experiment from the net absorbance on each subsequent day. The standardized area under the growth progress curve (sAUGPC) was calculated using the trapezoidal integration method standardized by duration of the experiment in days (Simko & Piepho, 2012).

For SP samples, net polyamine content was calculated by subtracting the mean concentration for each polyamine measured of the non-inoculated negative control medium (PD3). This correction was not applied to the pellet samples, as centrifugation removes the influence of the initial medium. Any negative values after normalization were set to zero.

Statistical analyses and data visualization were conducted using R software. Given the variability of bacterial growth and polyamines production, Wilcoxon rank-sum tests were used for all pairwise comparisons (*P* ≤ 0.05). For minimal media and biofilm experiments, each tested medium was compared against the PD3 control, and between media pairs sharing the same carbon source. For polyamine data, differences between time points within each sample type and strain were assessed.

Data were visualized using violin plots for growth and biofilm formation in minimal media, and polyamine data. For polyamine results, a pseudo-logarithmic scale was used to facilitate comparison across several orders of magnitude. All visualizations were produced using ggplot2, ggpubr and ggh4x packages in R.

## 3. Results

### 3.1. Metabolic characteristics of the *Xylella fastidiosa* pangenome and the core metabolic model

A curated genome-scale metabolic reconstruction was generated based on the core genes of 18 *X. fastidiosa* strains encompassing five subspecies (Table 1), including the strains previously used for metabolic modelling (Gerlin et al., 2020; Oliveira et al., 2023). Host diversity of the selected strains reflects the broad ecological range of the species and provides a representative basis for assessing the pangenomic and metabolic variability of this phytopathogen. Moreover, this pangenome also incorporates recently emerged European strains, with six isolates from Spain, Italy and France, in order to account for the lineages involved in the outbreaks starting to be detected in 2013, which have resulted in substantial agricultural and economic losses in Europe (Sanchez et al., 2019; De la Fuente, 2024).

Pangenome analysis revealed a total of 2660 ortholog groups (OGs), comprising 40828 genes, of which 1372 OGs included genes found in the 18 strains (Fig. 1A). The core genome consists of 26831 genes, constituting 65.72% of the genes analyzed. The accessory genome includes 13421 genes, corresponding to 32.87% of the total pangenome. Among the analyzed strains, XYL468 exhibited the highest number of accessory genes (890), whereas Salento-1 displayed the lowest (595). A total of 576 genes were classified as unique, accounting for 1.41% of all genes. The strain with the largest number of unique genes was XYL1961/18, with 141 unique genes, while U24D contained only 2 unique genes. Only a limited number of OGs showed strict subspecies-specific distribution.

**Figure 1.**
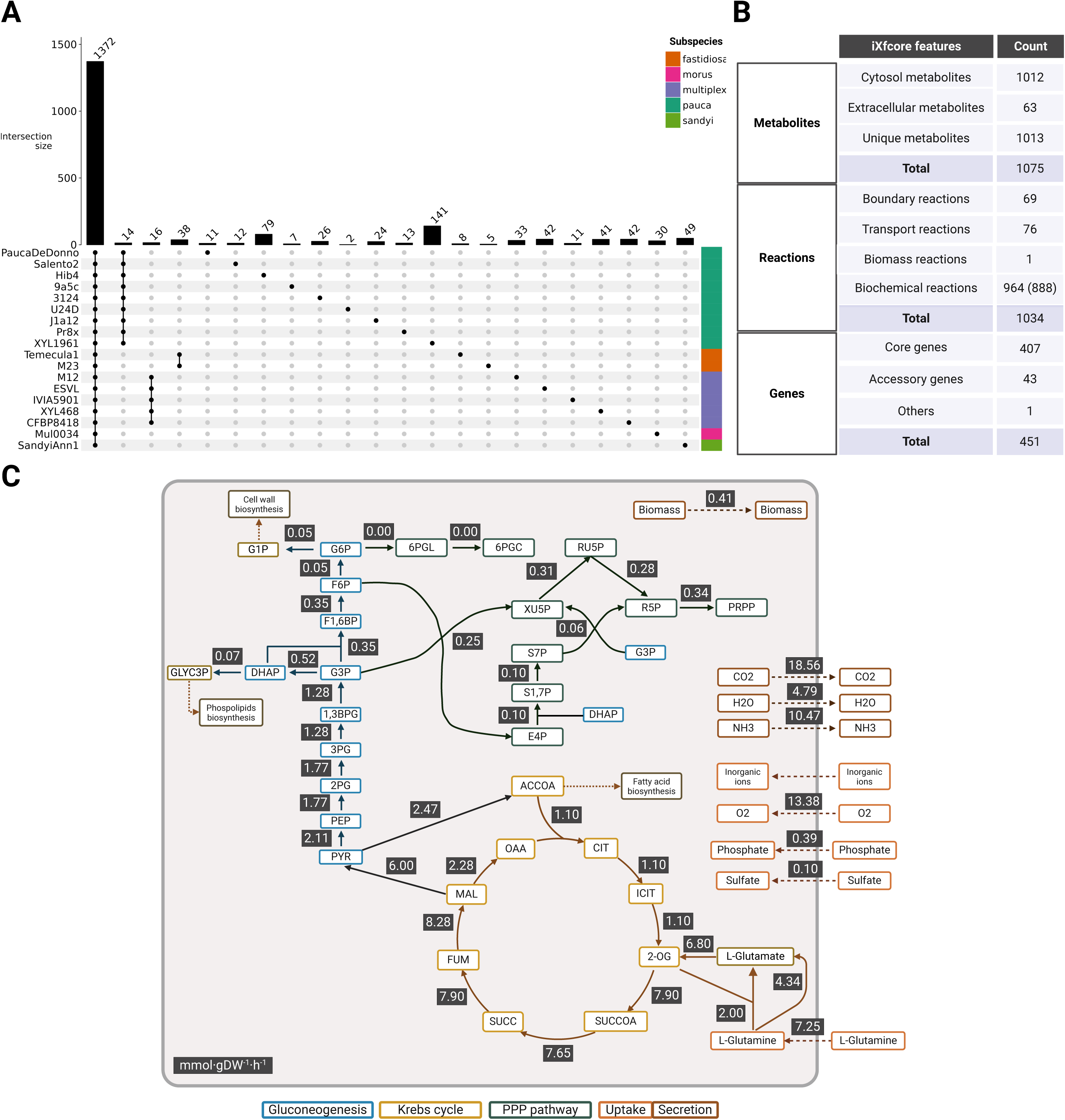
Genome-scale metabolic model reconstruction of *Xylella fastidiosa* through a pangenome based on the subspecies *pauca*, *multiplex*, *fastidiosa*, *morus* and *sandyi*. A) Pangenome analysis illustrating the distribution of core genes, genes shared between subspecies and unique genes. Each bar in the vertical bar chart represents the number of ortholog groups shared by the strains indicated below the chart. B) Summary of the features (reactions, metabolites and genes) from the *X. fastidiosa* metabolic core (iXfcore) reconstructed in this work. For further information on accessory genes, see Supplementary Results and Discussion, and Table S2. C) Predicted flux distribution through the core metabolism, using FBA, in a minimal medium with glutamine as the sole carbon and nitrogen source, achieving a specific growth rate of 0.41 h^-1^. Metabolic fluxes (mmol·gDW⁻¹·h⁻¹) are portrayed numerically.

Functional annotation of the pangenome revealed clear differences between the core, accessory, and unique gene sets (Fig. S1). Core OGs were predominantly enriched in metabolic functions, particularly carbohydrate, amino acid, nucleotide, energy and glycan metabolism, as well as metabolism of cofactors and vitamins. These were followed in prevalence by genes associated with genetic information processing and with signalling and cellular processes. In contrast, accessory genes showed a stronger representation of the latter two categories, including a higher proportion of functions associated with prokaryotic defense, secretion and transport systems, and quorum-sensing pathways, alongside a substantial fraction of genes with unknown function. Metabolic genes were also present within the accessory set, although to a lesser extent. For the unique gene set, only 6.2% (36 out of 576) of the genes could be functionally annotated, and none were linked to core metabolic pathways. Instead, the annotated genes mapped to categories such as prokaryotic defense systems, quorum sensing, or DNA replication and repair.

The core ortholog set identified in the pangenome was used to generate a draft genome-scale metabolic reconstruction using the automated reconstruction software ModelSEED (Henry et al., 2010). Curation was performed iteratively (see Supplementary Results and Discussion). The features of the final genome-scale metabolic reconstruction (iXfcore; Supplementary Material S1) are summarized in Fig. 1B. From a total of 1034 reactions, 83% are associated with gene-protein-reaction (GPR) rules. Of the reactions lacking GPR associations, 57% correspond to boundary or transport reactions, the majority of which do not require gene assignment. Model quality and consistency evaluated using the Memote testing suite (Lieven et al., 2020), yielded a total score of 85% and a consistency score of 97% (Supplementary Material S2). Annotation completeness scores were 66% for metabolites, 73% for reactions, and 53% for genes. The coverage of SBO terms reached 83%, indicating strong semantic and structural annotation.

The core gene set of *X. fastidiosa* encoded almost all central metabolic functions, although several specific gaps and distinctive features were identified (see Supplementary Results and Discussion). Of particular relevance is the absence in all analyzed strains of a glyoxylate cycle, as well as the lack of anaplerotic reactions connecting the intermediates of glycolysis and the TCA cycle.

Finally, the biomass reaction was formulated to include all known cellular components and their respective contributions per gram of biomass. Following extensive curation, including comparative evaluation against the biomass formulations of the *E. coli* model iJO1366 (Orth et al., 2011) and the *X. fastidiosa* model Xfm1158 (Gerlin et al., 2020), the biomass equation included 78 metabolites.

FBA predicted a maximum specific growth rate of 0.413 h⁻¹ when glutamine, one of the major components of xylem sap, was supplied as the sole carbon source at an uptake rate of 7.25 mmol·gDW⁻¹·h⁻¹ (Supplementary Material S3). The corresponding optimal flux distribution is shown in Fig. 1C. Under these conditions, most of the imported glutamine is converted into α-ketoglutarate. Carbon flux proceeds through the TCA cycle and malate is partially converted to pyruvate via the malic enzyme, generating NADPH. Pyruvate bifurcates toward gluconeogenesis or to the synthesis of acetyl-CoA. The iXfcore model further predicted uptake of inorganic ions, phosphate, and sulfate under these growth conditions, while CO₂ and NH₃ were secreted as metabolic by-products. Oxygen was also required, as the model did not predict growth under anaerobic conditions.

### 3.2. Growing *Xylella fastidiosa* in minimal media predicted by the model

Empirical efforts have led to the development of several culture media for *X. fastidiosa*, including PD2 (Davis et al., 1980), BCYE (Wells et al., 1981) and PW (Davis et al., 1981). However, these formulations do not fully overcome the fastidious growth requirements of the bacterium. In this work, a reverse ecology approach based on constraint-based metabolic modeling was applied to predict a synthetic defined minimal medium capable of sustaining biomass production in the iXfcore model. Under a fixed L-glutamine uptake constraint (7.25 mmol·gDW⁻¹·h⁻¹), the minimal medium prediction identified oxygen, phosphate, sulfate, glutamine (as both carbon and nitrogen source), and inorganic ions as required components (see Supplementary Material S4 for the complete list of metabolites and corresponding uptake fluxes).

Based on these predicted requirements, six synthetic defined minimal media were formulated (mXC1–mXC6; Table S1). Glutamine was used as the sole carbon and nitrogen source in mXC1 and mXC2, given its abundance in xylem sap. Two different iron sources were tested in these media, ferric pyrophosphate and hemin chloride, to assess iron bioavailability. Acetate was used as the carbon source in mXC3 and mXC4, supplemented with ammonium sulfate as a nitrogen source. To validate model predictions regarding biotin dependency under acetate growth conditions (see section 3.3), mXC4 was additionally supplemented with biotin. Finally, ribose was tested as the sole carbon source in mXC5 and mXC6. In mXC6, cysteine was evaluated as an alternative nitrogen source, as it showed strong utilization in nitrogen assimilation assays (data not shown).

All minimal media supported, to some extent, the growth and biofilm formation in all *X. fastidiosa* strains studied (Fig. 2; Supplementary Material S4). Interestingly, all strains were able to grow with glutamine media, despite glutamine not supporting growth in the phenotypic characterization of Biolog PM plates (see Supplementary Results and Discussion). The reference medium PD3 significantly supported higher growth than all minimal media across strains (Fig. 2A), except for XYL1961/18, which showed higher growth in the Glutamine+Fe formulation (mXC1) and no significant differences were detected for the rest of the media. In strain XYL468, no significant differences were observed between PD3 and Glutamine+Hemin or Ribose+Cysteine media.

**Figure 2.**
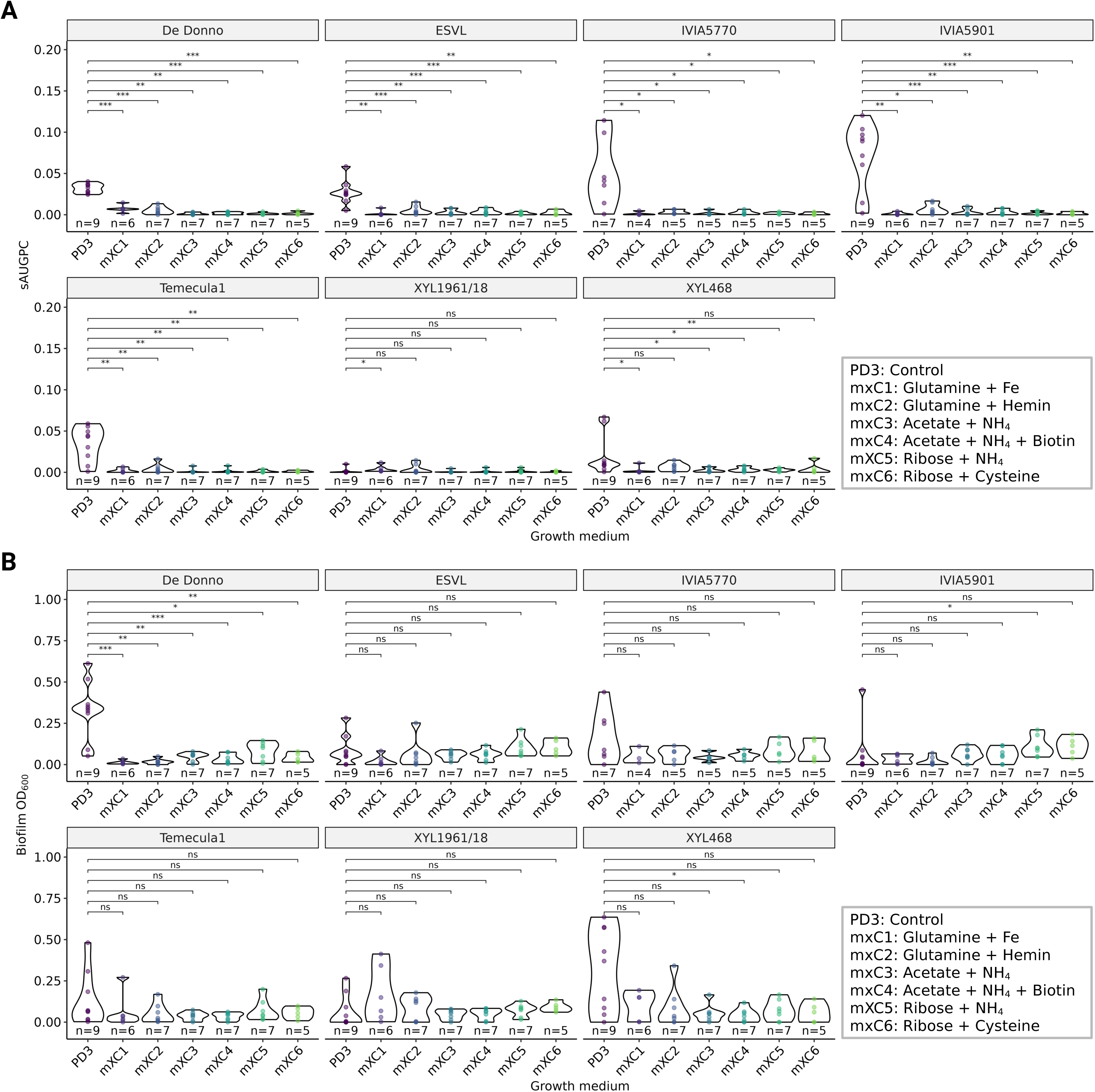
Evaluation of six synthetic defined growth media based on iXfcore model predictions. Total bacterial growth measured by the standardized area under the growth progress curve (sAUGPC) (A) and biofilm formation (B) of seven *Xylella fastidiosa* strains grow at 28 °C for 6 days. PD3 medium is included as a positive control. Statistical differences between each synthetic defined medium and the PD3 control were assessed using Wilcoxon rank-sum tests. Significance levels of comparisons are displayed above each line connecting group pairs (ns: *P*>0.05, ∗: *P*≤0.05, ∗∗: *P*≤0.01, ∗∗∗: *P*≤0.001, ∗∗∗∗: *P*≤0.0001).

Biofilm formation (Fig. 2B) showed comparable levels across most minimal media relative to PD3. The exception was strain De Donno, which produced significantly higher biofilm in the control medium PD3. Overall, a general trend was observed in which synthetic defined media containing ribose supported higher biofilm production across most strains, followed by acetate-based media (Fig. 2B).

Pairwise comparisons between media formulations sharing the same carbon source revealed no significant differences for growth or biofilm formation associated with iron source, biotin supplementation in acetate media, or nitrogen source substitution in ribose-based formulations (data not shown).

### 3.3. Acetate metabolism in *Xylella fastidiosa*

Experimental evidence supports the ability of *X. fastidiosa* to grow on acetate as the sole carbon source in several strains (Gerlin et al., 2020; this work, see Supplementary Results and Discussion). Here, all previously described metabolic solutions to produce biomass from acetate were evaluated, namely: (i) the glyoxylate cycle (Kornberg & Krebs, 1957); (ii) a ferredoxin-dependent pyruvate synthase (Segura et al., 2008), typically associated with anaerobic metabolism; (iii) the ethylmalonyl-CoA (EMC) pathway (Erb et al., 2007); (iv) the methylaspartate cycle (MaC) characteristic of haloarchaea (Khomyakova et al., 2011); and (v) part of the 3-hydroxypropionate (3HP) autotrophic bi-cycle (Herter et al., 2002). Across the 18 strains comprising the pangenome, none of these previously described pathways were fully supported by genomic evidence, lacking all or some of the necessary enzymes.

The iXfcore metabolic model predicted growth on acetate through an alternative, yet undescribed metabolic solution (Fig. 3). This consisted of a modular pathway with 11 steps (Table S3), connecting acetate to central carbon metabolism, generating pyruvate and succinate and enabling net carbon assimilation into biomass. This pathway integrates two enzymatic subsets present in previously described metabolic routes. The first one corresponds to the initial half of the 3HP bi-cycle, linking acetyl CoA to the synthesis of propionyl-CoA (the remaining reactions of the cycle are absent from the pangenome). The second subset corresponds to the methylcitrate pathway, which enables the assimilation of propionyl-CoA into pyruvate and succinate (Brämer & Steinbüchel, 2001; Dolan et al., 2018). During the reconstruction process, minor gaps were identified in the draft network. These gaps were resolved through manual genome evaluation and literature reviewing. All reactions included in the final pathway are supported by genes present in the pangenome. The putative acetate assimilation pathway in *X. fastidiosa* is represented in Fig. 3 and the Supplementary Results and Discussion file contains a detailed explanation of it.

**Figure 3.**
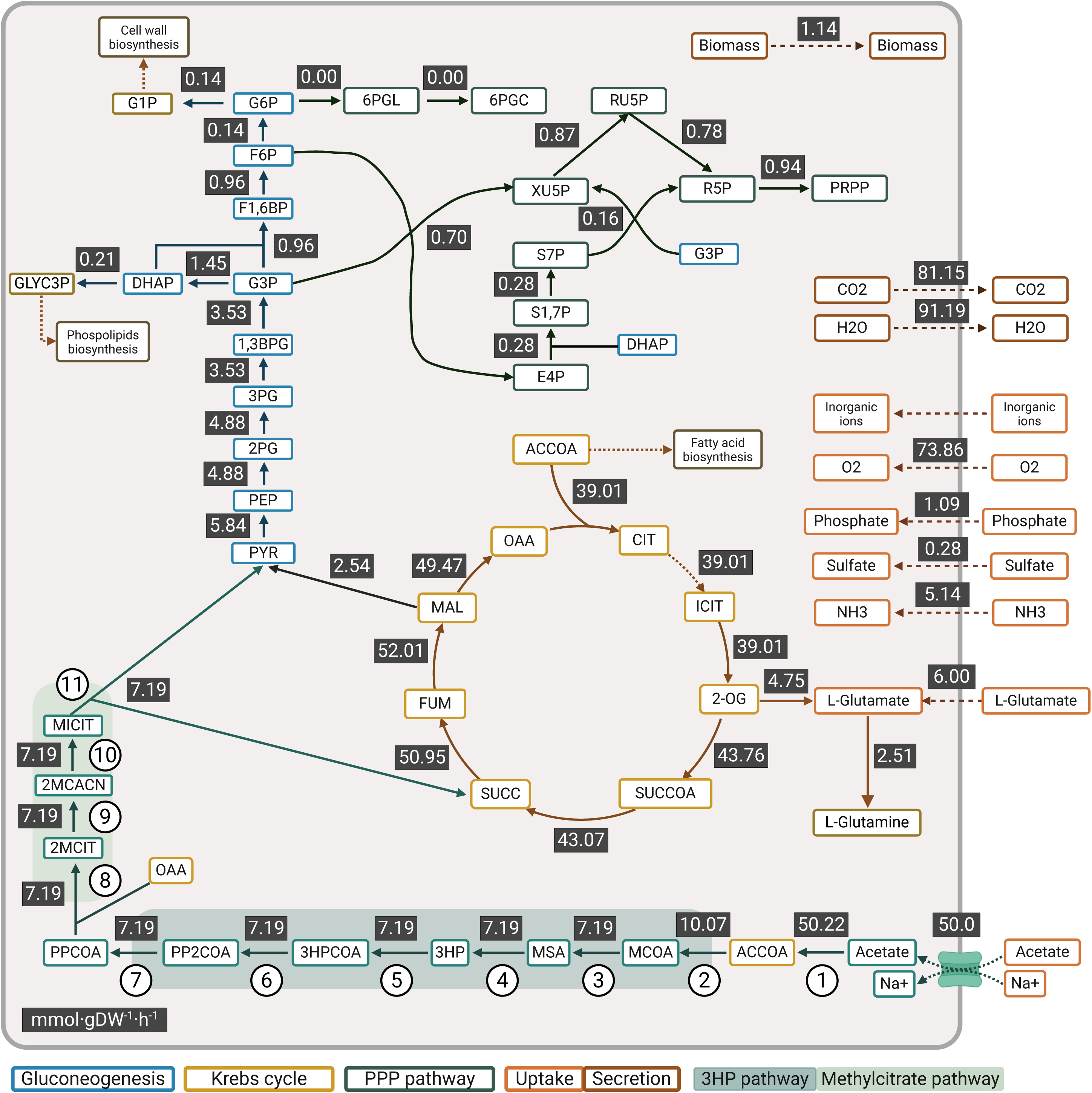
Predicted acetate assimilation by the iXfcore model. Flux distribution through the *Xylella fastidiosa* core metabolism, using FBA, in medium with acetate as the sole carbon and glutamic acid and ammonium as nitrogen source, achieving a specific growth rate of 1.14 h^-1^. Metabolic fluxes (mmol·gDW⁻¹·h⁻¹) are portrayed numerically. The alternative metabolic strategy for converting acetate into biomass combines two previously described modules: half of the 3-hydroxypropionate (3HP) bicycle (Herter et al., 2002) and the methylcitrate pathway (Brämer & Steinbüchel, 2001; Dolan et al., 2018) for propionyl-CoA assimilation. The enzymes and genes involved in the metabolic steps (1-11) of this pathway are in Supplementary Table S4. Abbreviations for metabolites participating in the 3HP pathway and the methylcitrate pathway: MCOA (malonyl CoA), MSA (malonate semialdehyde or 3-oxopropanoate), 3HP (3-hydroxypropionate), 3HPCOA (3-hrydroxypropionyl CoA), PP2COA (acryloyl CoA), PPCOA (propionyl CoA), 2MCIT (2-methylcitrate), 2MCACN (2-methylaconitate), MICT (2-methylisocitrate).

Compared with the flux distribution obtained when glutamine was used as the sole carbon source (Fig. 1C), in the flux distribution corresponding to acetate assimilation (Fig. 3; Supplementary Material S5) pyruvate is no longer converted into acetyl-CoA, as this metabolite is directly generated from acetate (Fig. 3, step 1). The pyruvate generated in the proposed pathway is redirected toward gluconeogenesis, similar to the glutamine-based simulation. Under acetate conditions, the malic enzyme also contributes to NADPH generation, although at 42% of the flux of the one observed with glutamine. This reduction may seem a paradox since the acetate assimilation pathway proposed here shows an increased demand of reducing power. A higher proportion of flux is directed through oxaloacetate (95% vs 27.5% with glutamine). This increased TCA cycle flux implies a higher flux through isocitrate dehydrogenase (IDH). The iXfcore model contains both the NAD- and NADP-dependent IDH isozymes, and when the model was evaluated with pFBA under the same conditions, with acetate 76% of the flux through IDH was generating NADPH, whereas with glutamine as the only C source, all the flux went through the NAD-dependent IDH isoenzyme.

### 3.4. Polyamine production by *Xylella fastidiosa* strains

The iXfcore model includes the biosynthetic pathways for the polyamines putrescine, spermidine and spermine, including the S-adenosylmethionine (SAM) regeneration cycle required for spermidine and spermine synthesis (Fig. S2). In *R. solanacearum*, putrescine secretion has been experimentally reported (Lowe-Power et al., 2018), although no homologs of known polyamine exporters have been identified in its genome (Igarashi & Kashiwagi, 2010; Sugiyama et al., 2017), suggesting the existence of a yet uncharacterized transport system. Since *X. fastidiosa* lacks any genomic evidence supporting a polyamine export system, cytosolic demand reactions were introduced in the iXfcore model to simulate polyamine overproduction.

When polyamine demand reactions were enforced or used as the objective function (see section 2.10), the model predicted feasible production of putrescine, spermidine, and spermine, beyond biomass requirements (Fig. 4A). However, increasing polyamine production resulted in a progressive reduction in biomass flux, revealing a trade-off between growth and polyamine production. The extent of this trade-off varied among the three polyamines, with differences in maximum achievable fluxes at minimum biomass levels. When putrescine overproduction was constrained to zero, spermidine and spermine exhibited a linear inverse relationship, with limited flexibility concerning the production of biomass and combined polyamine (Fig. 4A). In contrast, when putrescine overproduction was allowed together with either spermidine or spermine, a broader feasible solution space for similar biomass levels was observed. In these cases, similar biomass levels could be obtained with different combinations of polyamine production.

**Figure 4.**
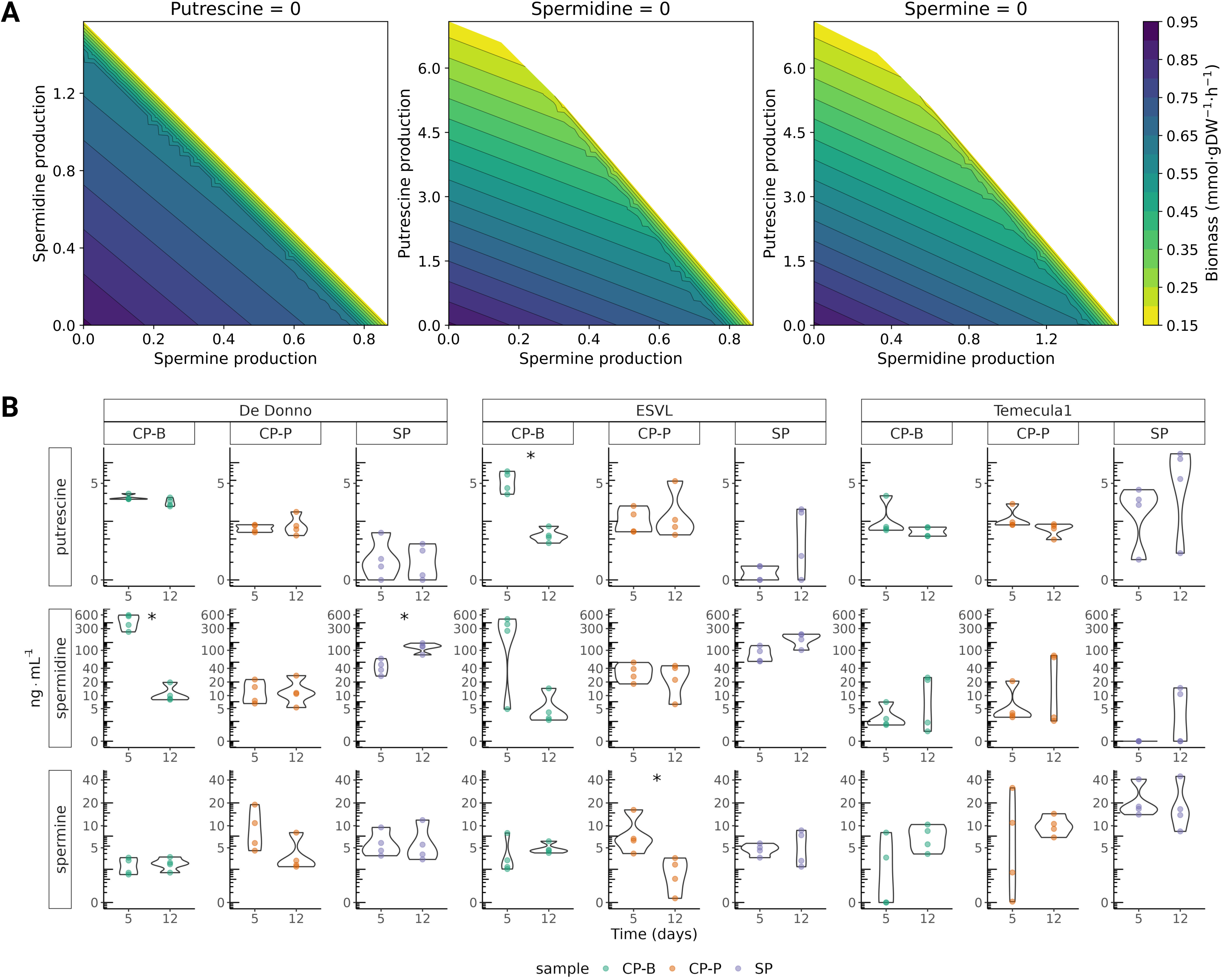
Polyamine production by *Xylella fastidiosa.* A) Trade-off between biomass and polyamine production using the iXfcore model. This figure illustrates the inverse relationship between growth (biomass accumulation) and the production of polyamines (putrescine, spermidine, and spermine) when one of the polyamines is completely absent. Simulations were performed using a minimal medium where glutamine served as the sole carbon source (with an uptake flux of 15.665 mmol·gDW⁻¹·h⁻¹). For each data point, the biomass reaction was maximized while the production fluxes for the polyamines (putrescine, spermidine, and spermine) were incrementally forced to the values represented by axis in flux units, mmol·gDW⁻¹·h⁻¹. A minimal threshold of 0.1608 mmol·gDW⁻¹·h⁻¹ was established for the consideration of growth. B) In vitro production of putrescine, spermidine and spermine across different sample types (cell pellet from biofilm, CP-B; cell pellet from planktonic growth, CP-P; and culture supernatant, SP). Samples were obtained from three *X. fastidiosa* strains (De Donno, ESVL and Temecula1) grown at 28 °C in PD3 medium for 5 and 12 days, with four replicates for each sample and time point. Concentrations (in ng·mL^-1^) are visualized with a pseudo-logarithmic scale. Statistical differences between time points within each sample type and strain were assessed using Wilcoxon rank-sum tests. Only significant results are indicated (∗: *P*≤0.05).

The experimental results on polyamine production by *X. fastidiosa* strains corroborated the findings of the iXfcore metabolic model (Fig. 4B and Fig. S3; Supplementary Material S6. In a first experiment six *X. fastidiosa* strains belonging to the three subspecies studied in this work produced and secreted polyamines in vitro (Fig. S3). Putrescine, spermidine and spermine were detected in both biofilm and planktonic growth, as well as in the culture supernatant. Spermidine was produced at higher levels than putrescine and spermine. In a second experiment, polyamine levels were analyzed in three distinct sample types: planktonic cell pellets, biofilm-forming cell pellets, and the culture supernatant. Again, spermidine was the most abundant polyamine detected (Fig. 4B). There were no significant differences observed between time points within each sample type, except for the four comparisons marked with an asterisk in Fig. 4B.

### 3.5. Trade-off between virulence factors and biomass

Previous reconstructions of *X. fastidiosa* metabolism (Gerlin et al., 2020; Oliveira et al., 2023) have considered the secretion of two known virulence factors: the LesA protein (Nascimento et al., 2016) and exopolysaccharides (EPS) (da Silva et al., 2001). The metabolic model of the phytopathogen *R. solanacearum* (Peyraud et al., 2016), also incorporated polyamine secretion. In this work, several scenarios were analyzed by simulating the combined production of LesA, EPS, and one of the detected polyamines: putrescine (Fig. 5A), spermidine (Fig. 5B), or spermine (Fig. 5C). Under these conditions, the model predicted maximum fluxes of 0.02 mmol·gDW⁻¹·h⁻¹ for LesA protein, 1.45 mmol·gDW⁻¹·h⁻¹ for EPS, 7.09 mmol·gDW⁻¹·h⁻¹ for putrescine, 1.57 mmol·gDW⁻¹·h⁻¹ for spermidine, and 0.87 mmol·gDW⁻¹·h⁻¹ for spermine.

**Figure 5.**
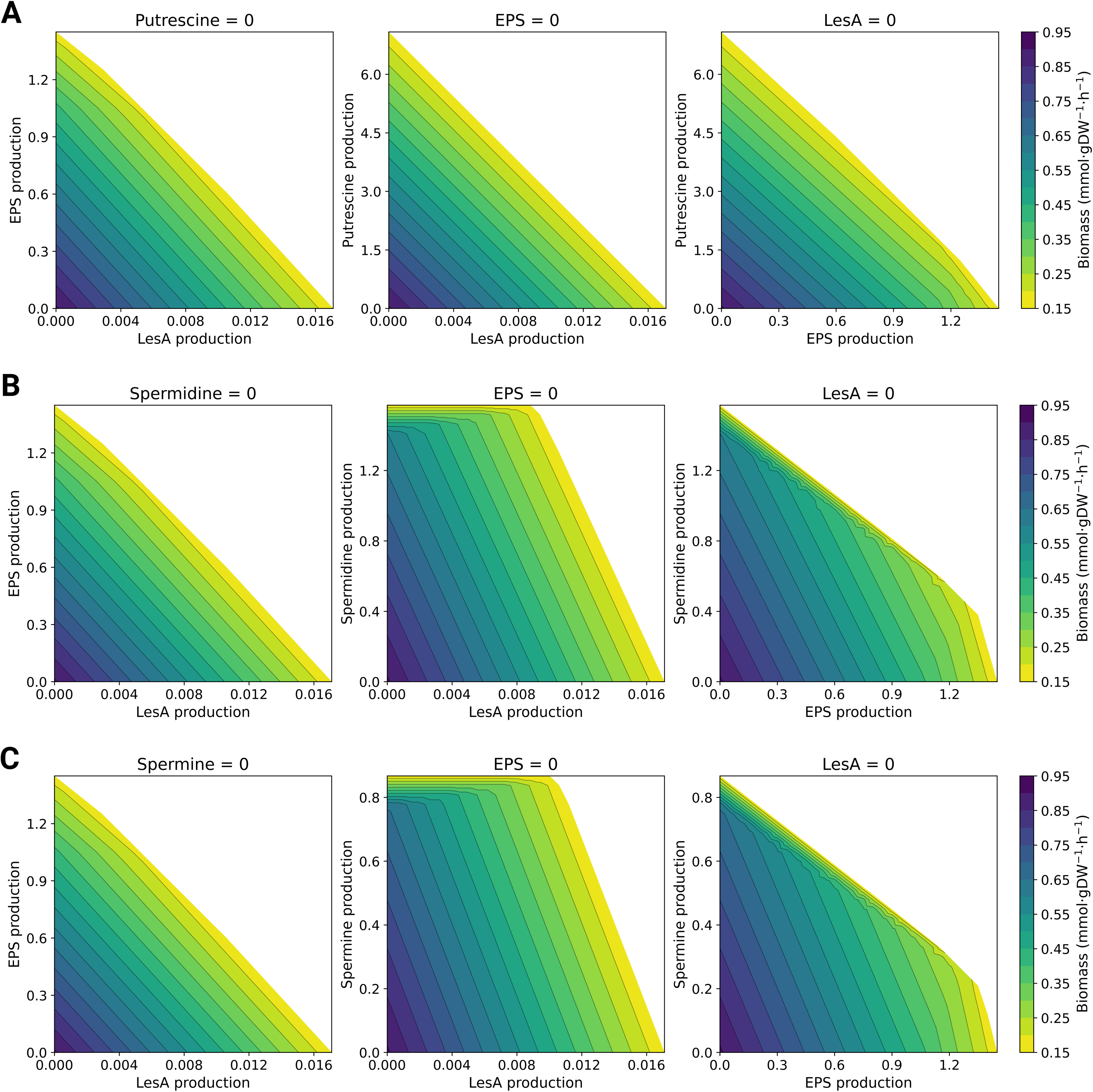
Trade-off between biomass and virulence factor production in *Xylella fastidiosa.* This figure illustrates the inverse relationship between growth (biomass accumulation) and the production of: LesA protein, fastidian gum EPS, and putrescine (A), spermidine (B) or spermine (C) using the iXfcore metabolic mode, when one of the three compounds is completely absent. Simulations were performed using a minimal medium where glutamine served as the sole carbon source (with an uptake flux of 15.664 mmol·gDW⁻¹·h⁻¹). For each data point, the biomass reaction was maximized while the production fluxes for the virulence factors (LesA, EPS, polyamine) were incrementally forced to the values represented by the axis in flux units, mmol·gDW⁻¹·h⁻¹. A minimal threshold of 0.1608 mmol·gDW⁻¹·h⁻¹ was established for the consideration of growth.

A clear trade-off between virulence factor secretion and biomass production was observed (Fig. 5). The combined production of LesA with EPS showed a linear inverse relationship. For the production of putrescine together with the other virulence factors (Fig. 5A), a nearly linear inverse relationship was observed between the combined production of these compounds and biomass formation. Interestingly, in the case of spermidine (Fig. 5B) and spermine (Fig. 5C), the feasible solution space showed greater compatibility between LesA production and polyamine secretion. In these scenarios, different rates of LesA production remain feasible even when spermidine or spermine are produced at high levels, indicating greater metabolic flexibility for LesA secretion while maintaining the same biomass flux. This behavior is reflected by the broader feasible region observed in the phase planes. In particular, a change in slope appears when LesA production reaches approximately 0.02 mmol·gDW⁻¹·h⁻¹. A similar but less pronounced pattern is observed for EPS production. Overall, EPS secretion appears to be more compatible with spermidine or spermine production than with putrescine production, particularly near the upper limit of EPS flux (around 1.3 mmol·gDW⁻¹·h⁻¹).

## 4. Discussion

In this work, we reconstructed the core metabolic model of *X. fastidiosa* based on the pangenome of 18 strains belonging to five subspecies of this phytopathogen. This approach accounts for the high genetic diversity within the species while allowing the identification of its conserved metabolic capabilities. Previous metabolic reconstructions of *X. fastidiosa* were strain-specific, including the model for the subsp. *multiplex* CFBP 8418 (Gerlin et al., 2020) and the model for the subsp. *pauca* De Donno (Oliveira et al., 2023). In the first study, genome comparisons were performed with the strains *X. fastidiosa* subsp. *pauca* 9a5c and subsp. *fastidiosa* Temecula1, suggesting that the results could be extrapolated to these subspecies. In contrast, our reconstruction explicitly incorporates these strains within a broader pangenome framework, providing a more robust representation of the conserved metabolic potential of *X. fastidiosa*.

Moreover, the iXfcore metabolic model and the pangenome assembly can serve as a foundation for the reconstruction of strain-specific models, following the protocol described in Norsigian et al. (2020). The availability of a curated-base model can significantly reduce the effort required for the generation of multi-strain metabolic models. Such approaches have already been successfully applied to reconstruct models for the *Lactobacillaceae* family and for *Aspergillus fumigatus* strains (Mirhakkak et al., 2023; Ardalani et al., 2024). Additionally, our reconstructed model incorporates the OG associated with each gene in the network annotation, enabling the extraction of strain-specific genetic information when required.

*X. fastidiosa* appears to lack typical anaplerotic reactions to replenish the TCA cycle, suggesting that network operation is dependent of an external supply of precursors of the cycle intermediates–such as the amino acids glutamine or glutamate available in the xylem. iXfcore model simulations using hexoses, pentoses or even acetate, are consistent with this observation: without such intermediates, the model cannot sustain growth. In previous reconstructions, different assumptions evaded this issue. Thus, the model by Gerlin et al. (2020) supported growth with the inclusion of the glyoxylate cycle despite without genetic evidence, whereas the reconstruction by Oliveira et al. (2023) allowed a reversible malic enzyme generating malate from pyruvate. If we constrain the malic enzyme to the oxidative decarboxylation direction, restauration of growth requires the presence of a TCA intermediate. In this case, the malic enzyme may contribute to NADPH production in the context of a network with a limited set of reactions for reducing power production. For instance, the in silico simulations indicated that the Entner–Doudoroff (ED) pathway represents the main route for sugar metabolism. This result is consistent with the results of Oliveira et al. (2023) that suggested that this could help protect the bacterium against reactive oxygen species (ROS) generated by the plant defense system (Wang et al., 2017). The generation of NADPH by the ED pathway has been associated with an increased tolerance to oxidative stress in *P. putida* (Chavarría et al., 2013).

In axenic cultures, *X. fastidiosa* can grow with acetate as a sole carbon source. Remarkably, acetate is a major organic acid present in olive xylem sap (Anguita-Maeso et al., 2021). Intriguingly, GEMs for this organism lack any canonical pathway for acetate assimilation, and previous metabolic reconstructions have addressed this issue using different strategies. As mentioned before, growth on acetate was enabled in the Xfm1158 model (Gerlin et al., 2020) due to the inclusion of a glyoxylate cycle without genomic evidence of its key enzymes. In contrast, Oliveira et al. (2023) adopted a more conservative position and the iMS50 model cannot simulate growth on acetate. In our reconstruction, the iXfcore model suggested a previously undescribed metabolic solution for acetate assimilation. In brief, we propose the first part of the 3HP bi-cycle–an autotrophic pathway described in *Chloroflexus auranticus* by Herter et al. (2002) (Fig. S4, panel A)–combined with the conversion of propionyl-CoA into a biomass precursor. The assimilation of propionyl-CoA in *X. fastidiosa* would use a pathway different to the one proposed by Zarzycki and Fuchs (2011) for the co-assimilation of acetate in *C. auranticus* (Fig. S4, panel B). Thus, in the iXfcore model propionyl-CoA can be assimilated into pyruvate through the methylcitrate pathway firstly described in yeasts by Tabuchi et al. (1974) and which is also operative in bacteria like *E. coli* grown in propionate (Brock et al., 2002) (Fig. S4, panel C).

We can compare this new pathway for acetate assimilation with the ones previously described in the literature (Table S4). The putative acetate assimilation pathway in *X. fastidiosa* (Xf pathway) represents a slight variation of the 3HP pathway described in *C. auranticus*, in terms of energy balance and carbon efficiency; in both cases, there is a net gain of carbon. The main difference is that the Xf pathway involves two more enzymatic steps than the half 3HP bi-cycle. Both pathways are more demanding of reducing power than the rest of known routes and, as discussed before, this may explain the increased flux through the TCA cycle in iXfcore under acetate conditions. It is remarkable that *X. fastidiosa* harbors two structurally different IDH isoenzymes, specific for NAD^+^ or NADP^+^, respectively (Lv et al., 2018). Interestingly, the evolutionary origin of a NADP-dependent IDH has been correlated with an increased demand of reducing power associated with the use of acetate as carbon source (Zhu et al., 2005).

Our proposed Xf pathway highlights the role of functional modularity in metabolic adaptation and evolution. Metabolic modules can reorganize or combine to explore biochemical diversity (Schada von Borzyskowski et al., 2020). Functional degeneracy between diverse metabolic solutions also can reflect different dynamics of adaptation to environmental challenges, as described for *Paracoccus denitrificans* (Kremer et al., 2019). On the other hand, in our hypothetical solution for acetate assimilation in *X. fastidiosa* we have proposed several steps catalyzed by enzymes with relaxed substrate specificity. Thus, an acyl-CoA synthetase operates in steps 1 and 5, a citrate synthase is also operative as methylcitrate synthase (step 8), and finally aconitase AcnD will act in step 10 (Fig. 3; Table S3; Supplementary Results and Discussion). Those cases would be examples of the well-known metabolic economy observed in symbionts (either pathogens or mutualists), sustained through reactions operated by promiscuous or multifunctional enzymes (Yus et al., 2009; Kim et al., 2010; Gil & Peretó, 2015). Gerlin et al. (2020) have correlated the lack of enzymatic redundancy in the *X. fastidiosa* metabolic network with a reduced robustness and its characteristic fastidious growth. In any case, we are aware that our in silico hypothesis requires experimental validation, through omics approaches, biochemical characterization of enzyme activities, and ^13^C-fluxomics experiments. Furthermore, phylogenetic analysis of the involved enzymes also could shed light on the origin and evolutionary tinkering of the pathway, as well as its distribution in the tree of life.

Metabolic reconstructions have been previously used to design minimal growth media for fastidious microorganisms. A notable example is the work for *Mycoplasma pneumoniae* to define a synthetic minimal medium (Yus et al., 2009). This strategy of reverse ecology implicitly validates that genomic information and reconstructed networks can guide the formulation of culture media for other fastidious or uncultured bacteria (Cross et al., 2019; Lewis et al., 2021). Under strictly defined conditions, the use of sugars or any other substrates that do not directly feed the TCA cycle would theoretically require the addition of a TCA intermediate. However, in empirical in vitro conditions, intracellular metabolite pools may partially compensate for this requirement, as cells are not metabolically empty at the beginning of the experiment. In natural environments, on the other hand, the chemical diversity accessible to the bacterium would compensate for this metabolic obstacle. Interestingly, minimal media tested in this study supported biofilm formation, with no significant differences compared to the standard media PD3. *X. fastidiosa* is known to colonize xylem vessels primarily in the form of biofilms, which play a key role in its pathogenicity (Marques et al., 2002). This approach aimed to explore whether minimal environmental conditions, such as in the xylem sap, could trigger biofilm development independently of an optimal planktonic growth.

One of the most notable findings of this work is the first direct experimental evidence of polyamine production in *X. fastidiosa*. Polyamines are well-established modulators of biofilm formation and adhesion in many bacterial pathogens. They influence biofilm development through effects on gene regulation, signaling pathways, and extracellular matrix production. However, until now there was no direct experimental evidence of polyamines production by *X. fastidiosa*. A previous study in citrus trees infected with this phytopathogen reported high levels of putrescine (Purcino et al., 2007), although this observation was interpreted as a stress signalling response by the plant host (Blázquez, 2024). In other phytopathogens, such as *R. solanacearum* and *P. syringae,* polyamines have been considered as a virulence factor (Lowe-Power et al., 2018; Solmi et al., 2025). In these pathogens, polyamines likely protect bacteria against oxidative stress generated by plant defense responses and may contribute to symptom development during infection.

Metabolic modelling has previously been used to explore production of virulence-associated processes in plant pathogens. Other reconstructions of *X. fastidiosa* metabolism (Gerlin et al., 2020; Oliveira et al., 2023) considered the secretion of two known virulence factors: the LesA protein (Nascimento et al., 2016) and exopolysaccharides (EPS) (da Silva et al., 2001). The metabolic model of *R. solanacearum* (Peyraud et al., 2016), also included polyamine secretion. We extended this concept by exploring the relationship between polyamine production and other virulence-associated factors in *X. fastidiosa*, namely LesA and EPS.

In this work, both metabolic modelling and experimental analyses indicated that *X. fastidiosa* is capable of producing and secreting polyamines. Our simulations revealed a trade-off between biomass growth, polyamine biosynthesis, and the production of other virulence factors, suggesting that the synthesis of these compounds is metabolically constrained in this bacterium. Spermidine and spermine production appear to be metabolically coupled due to their dependence on shared precursors. Additionally, the simulations revealed a trade-off between biomass production and the secretion of virulence factors. For instance, spermidine and spermine production appeared to be more compatible with the secretion of LesA and EPS across a broader range of similar biomass levels.

Our experimental data, despite the variability typically observed when working with fastidious bacteria in vitro, showed a higher accumulation of spermidine in the culture supernatant. Additionally, polyamine levels tended to be higher in biofilms than in planktonic cells, suggesting a link between polyamine production and the biofilm state of this phytopathogen. While polyamines are established regulators of bacterial biofilms (Karatan & Michael, 2013) in *X. fastidiosa* this role remains untested experimentally. Current investigations in *X. fastidiosa* focus on EPS composition or processing (Janissen et al., 2015; Castro et al., 2023), the role of surface adhesins and pili (Scala et al., 2025), or of DSF-family quorum sensing signals (Horgan et al., 2024), without an explicit link to polyamine metabolism or signalling. Our results open the possibility that polyamines could represent an additional virulence factor in *X. fastidiosa*. Further in vivo studies will be necessary to determine to which extension bacterial metabolism is responsible of the polyamines detected in infected plants, and to clarify their role during host colonization by this species.

## 5. Conclusions

This work presents the first pangenome-based metabolic reconstruction for *X. fastidiosa*, integrating the conserved metabolic features shared across five subspecies of this plant pathogen. Following experimental validation, the iXfcore model successfully predicted minimal nutritional requirements, which were then used to design defined media capable of supporting biofilm formation in vitro. This outcome provided additional support for the model’s predictive accuracy. The model also revealed a previously undescribed pathway for incorporating acetate into biomass precursors. In addition, simulations predicted that *X. fastidiosa* can overproduce polyamines—a finding that was later confirmed experimentally, providing the first in vitro evidence of polyamine production in this pathogen. Beyond these specific findings, the iXfcore model establishes a robust systems-level framework for building strain-specific metabolic models. This framework can be used to further explore the metabolism of *X. fastidiosa*, generate new hypotheses about its physiology and virulence mechanisms, and support future studies that integrate bacterial and plant metabolism to better understand host–pathogen interactions.

## Supporting information

Supplementary Results and Discussion

## Funding sources

P.C.-A. was recipient of a predoctoral fellowship from the Generalitat Valenciana “Subvenciones para la contratación de personal investigador de carácter predoctoral (ACIF)”, with reference ACIF/2021/110. M.Á.-H. was supported by the Generalitat Valenciana and the European Social Fund “ESF Investing in your future” through grant CIACIF/2022/333. J.P. acknowledges the generic support of CYTED (Iber-Xyfas network “Red Iberoamericana para la vigilancia de *Xylella fastidiosa*”, ref. 119RT0569) as well as of his host institutions Universitat de València and CSIC. This work was financed by the HORIZON-CL6-2021-FARM2FORK-01 project BeXyl (Beyond *Xylella*, Integrated Management Strategies for Mitigating *Xylella fastidiosa* impact in Europe; Grant ID No 101060593). The funders had no role in study design, data collection and analysis, decision to publish, or presentation of the manuscript.

## CRediT authorship contribution statement

**Paola Corbín Agustí**: Conceptualization, Methodology, Formal analysis, Visualization, Investigation, Software, Data curation, Writing– Original draft, Reviewing and Editing. **Miguel Álvarez Herrera**: Methodology, Formal analysis, Visualization, Investigation, Software, Writing– Reviewing and editing. **Miguel Román Écija**: Methodology, Formal analysis, Investigation, Writing– Reviewing and editing. **Patricia Álvarez**: Methodology, Investigation, Software, Writing– Reviewing and editing. **Marta Tortajada**: Conceptualization, Writing–Reviewing and editing. **Juli Peretó**: Conceptualization, Supervision, Funding acquisition, Writing– Original draft, Reviewing and Editing. **Blanca B. Landa**: Conceptualization, Supervision, Funding acquisition, Writing– Original draft, Reviewing and Editing.

## Declaration of Competing Interest

M. Tortajada is a former employee of Biopolis-Archer Daniels Midland and contributed to this study in their personal capacity. The authors declare no conflicts of interest.

## Data Availability

The code and data for reproducing this analysis are available in the Zenodo repository: https://doi.org/10.5281/zenodo.17307291

## Acknowledgements

The authors thank the expert support of Miguel Ángel Blázquez and Francisco Vera-Sirera (IBMCP, UPV-CSIC) and the IBMCP Metabolomics Facility for the polyamine analyses. The authors thank Gonzalo de Oyanguren for the initial steps into the pangenome analysis. Figures 1C and 3 were created in BioRender. Smith, J. (2025).

## Appendix A. Supplementary material

### Supplementary Figures and Tables

PDF document containing supplementary figures and tables referred in the main text.

### Supplementary Methods

PDF document describing metabolic model refinement and curation, additional in silico constraints and simulation conditions, and phenotypic characterization of *X. fastidiosa* strains.

### Supplementary Results and Discussion

PDF document describing additional metabolic features of the iXfcore model, phenotypic characterization of *X. fastidiosa* strains, and the proposed pathway for acetate assimilation.

### Supplementary Material 1 (SM1). iXfcore metabolic model

Compressed archive (.zip) containing the model in SBML (.xml), JSON (.json), and MATLAB (.mat) formats.

### Supplementary Material 2 (SM2). MEMOTE report for iXfcore

HTML report detailing the results of the memote test suite, including network consistency, annotation coverage for metabolites, reactions, genes, and SBO terms and total score.

### Supplementary Material 3 (SM3). Flux distribution in minimal medium with glutamine as carbon and nitrogen source

Excel file reporting FBA and pFBA flux distributions (mmol·gDW⁻¹·h⁻¹) across reactions under the simulated minimal medium conditions.

### Supplementary Material 4 (SM4). Defined minimal media results

Excel file including in silico minimal media prediction and experimental measurements of growth and biofilm formation across the six designed synthetic minimal media.

### Supplementary Material 5 (SM5). Flux distribution under acetate as sole carbon source

Excel file reporting FBA and pFBA flux distributions (mmol·gDW⁻¹·h⁻¹) across reactions under acetate growth conditions.

### Supplementary Material 6 (SM6). Polyamine experimental data

Excel file containing experimental quantification of putrescine, spermidine, and spermine across the two different experiments.

### Supplementary Material 7 (SM7). Phenotypic Biolog growth data

Excel file containing PM1 and PM2 sAUGPC values, normalized data, and growth classification across the tested carbon sources.

### Supplementary Material 8 (SM8). In silico growth predictions

Excel file reporting FBA-predicted optimal growth fluxes (mmol·gDW⁻¹·h⁻¹) for the evaluated carbon sources.

## Notes

https://doi.org/10.5281/zenodo.17307291

### Competing Interest Statement

The authors have declared no competing interest.

### Summary of Updates

Main text revised and synthesized for clarity; Figures revised and updated; Supplementary material, data and code added.

