## Supplementary Figures and Tables for "A metabolic model based on a pangenome core unveils new biochemical features of the phytopathogen *Xylella fastidiosa*"

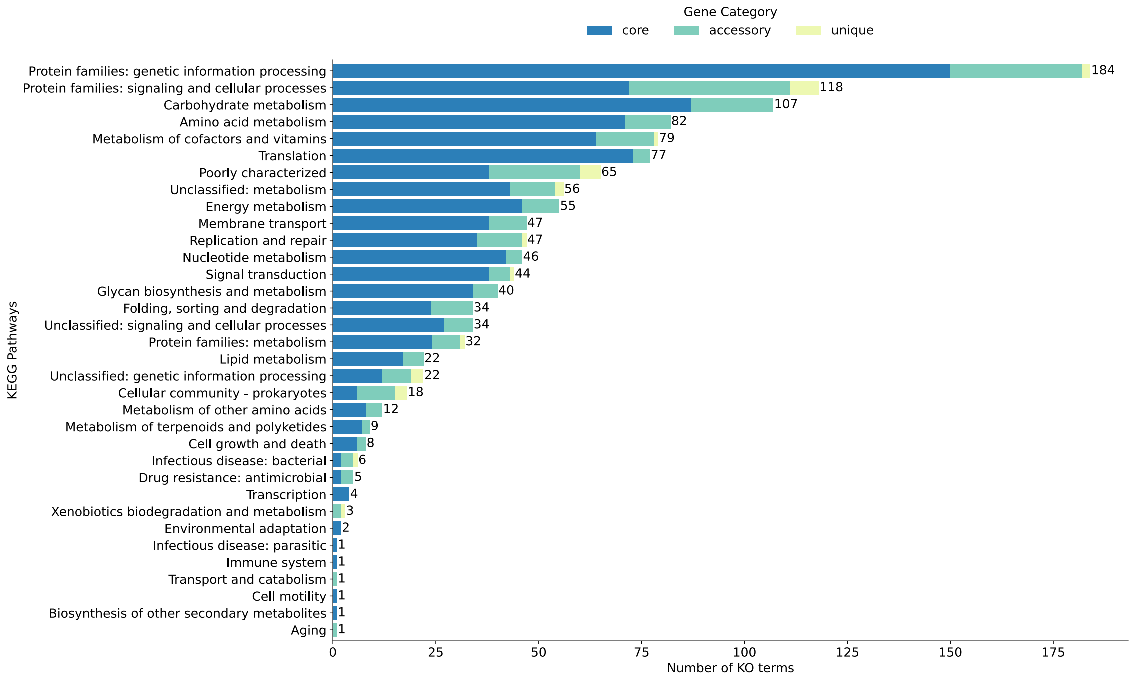


**Figure S1. Functional annotation of the core, accessory, and unique gene sets of the *Xylella fastidiosa* pangenome.** Functional categories were assigned based on KEGG Orthology (KO) annotation. The distributions represent the number of unique KO terms within Level 2 functional classifications of the KEGG hierarchy.

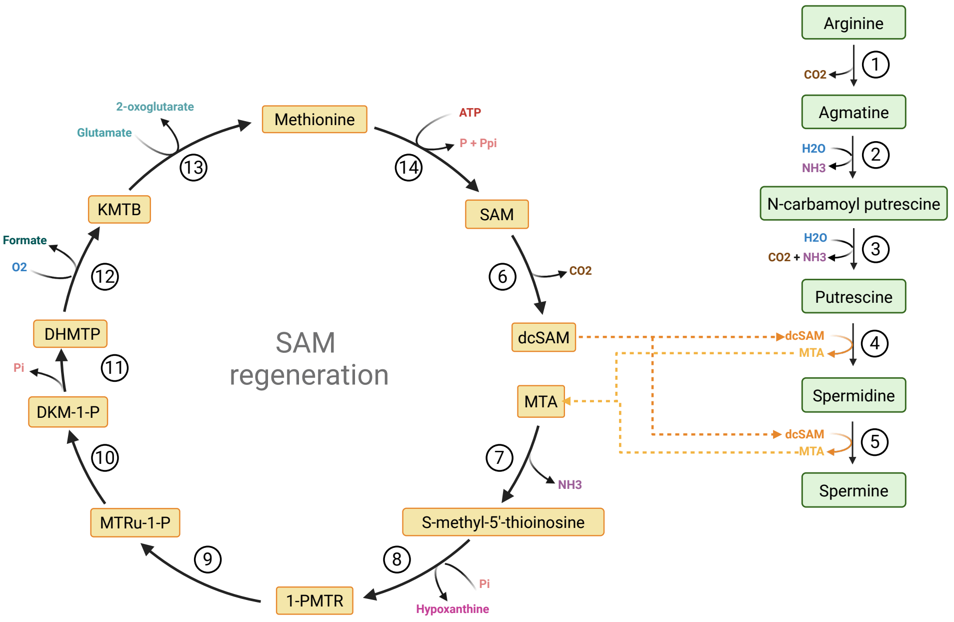


| Step | EC number | Enzyme | Locus tag | Reaction ID |
|---|---|---|---|---|
| 1 | 4.1.1.19 | Arginine decarboxylase | XF_RS00600 | rxn00405 |
| 2 | 3.5.3.12 | Agmatine deiminase | XF_RS10565 | rxn01029 |
| 3 | 3.5.1.53 | N-carbamoylputrescine amidase | XF_RS10570 | rxn00853 |
| 4 | 2.5.1.16 | Spermidine synthase | XF_RS00595 | rxn01406 |
| 5 | 2.5.1.22 | Spermine synthase | XF_RS00595 | rxn02061 |
| 6 | 4.1.1.50 | Adenosylmethionine decarboxylase | XF_RS06515 | rxn00127 |
| 7 | 3.5.4.31 | S-methyl-5'-thioadenosine deaminase | XF_RS10700 | rxn16503 |
| 8 | 2.4.2.44 | S-methyl-5'-thioinosine phosphorylase | XF_RS10205 | rxn16511 |
| 9 | 5.3.1.23 | S-methyl-5-thioribose-1-phosphate isomerase | XF_RS11120 | rxn03057 |
| 10 | 4.2.1.109 | Methylthioribulose 1-phosphate dehydratase | XF_RS09610 | rxn05104 |
| 11 | 3.1.3.77 | Acireductone synthase | XF_RS09620 | rxn05107 |
| 12 | 1.13.11.54 | Acireductone dioxygenase | XF_RS09615 | rxn05092 |
| 13 | 2.6.1.57 | Aromatic-amino-acid transaminase | XF_RS00150 | rxn05108 |
| 14 | 2.5.1.6 | Methionine adenosyltransferase | XF_RS01640 | rxn00126 |

**Figure S2. Polyamine biosynthesis pathway in *Xylella fastidiosa*.** Schematic representation of the biosynthetic pathway for putrescine, spermidine and spermine from arginine, including the S-adenosylmethionine (SAM) regeneration cycle required for spermidine and spermine synthesis in the iXfcore metabolic model. For each enzymatic step shown (1–14), the corresponding enzyme, EC number, associated gene(s) identified in *X. fastidiosa*, and the reaction ID in the iXfcore model are provided.

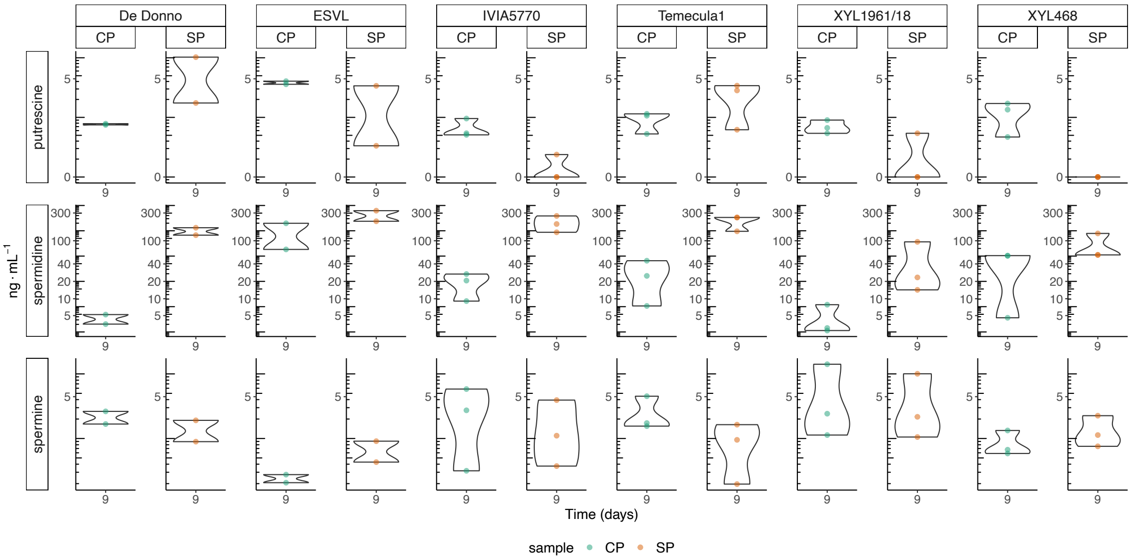


**Figure S3. In vitro production of putrescine, spermidine and spermine by *Xylella fastidiosa*.** Cell pellet from biofilm and planktonic growth (CP) and culture supernatant (SP) samples were obtained from six *X. fastidiosa* strains (De Donno, ESVL, IVIA5770, Temecula1, XYL1961/18 and XYL468) grown at 28°C in PD3 medium for 9 days, with at least two replicates for each sample. Concentrations (in ng·mL$^{-1}$) are visualized with a pseudo-logarithmic scale.

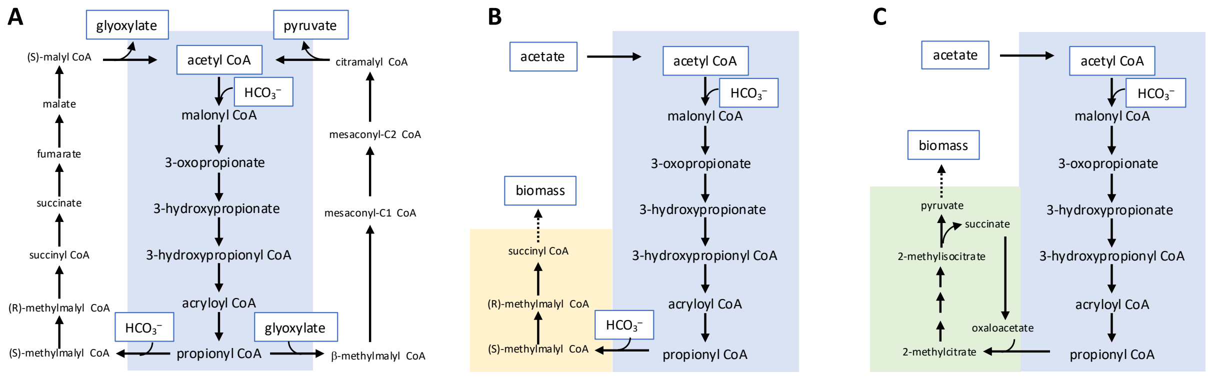


**Figure S4**. **Acetate assimilation based on part of the 3-hydroxypropionate (3HP) bi-cyle**. (A) The autotrophic 3HP bi-cycle as described in *C. auranticus*. The part of the bi-cycle from acetyl CoA to propionyl CoA is highlighted in blue. (B) *C. auranticus* can co-assimilate acetate using the central part of the 3HP bi-cycle (blue) with some additional steps allowing the transformation of propionyl CoA into a biomass precursor like succinyl CoA (yellow). (C) As described in this work, in *X. fastidiosa* the central module of the 3HP bi-cycle is combined with an alternative way to transform propionyl CoA into a biomass precursor (namely, pyruvate) using the methylcitrate pathway (green). For further details on (A) and (B) see Schada von Borzyskowski et al. (2020).

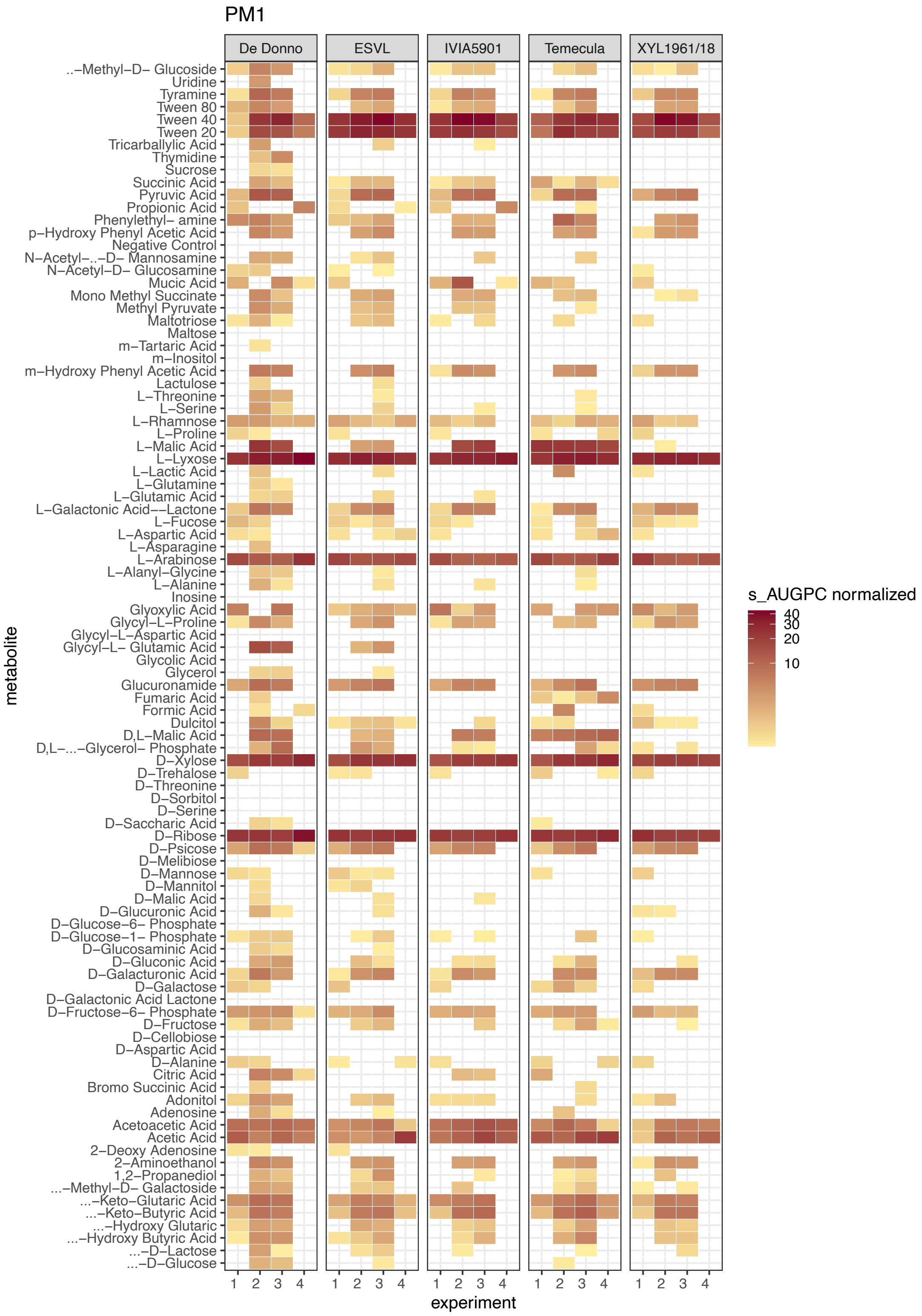


**Figure S5. Bacterial growth in the Biolog Phenotype MicroArray plates PM1 for five *Xylella fastidiosa* strains** (De Donno, ESVL, IVIA5901, Temecula1, and XYL1961/18) used in this study. Four independent experiments were performed for each strain. Growth is presented as the normalized standardized area under the growth curve (sAUGPC). Data is visualized using a pseudo-logarithmic scale.

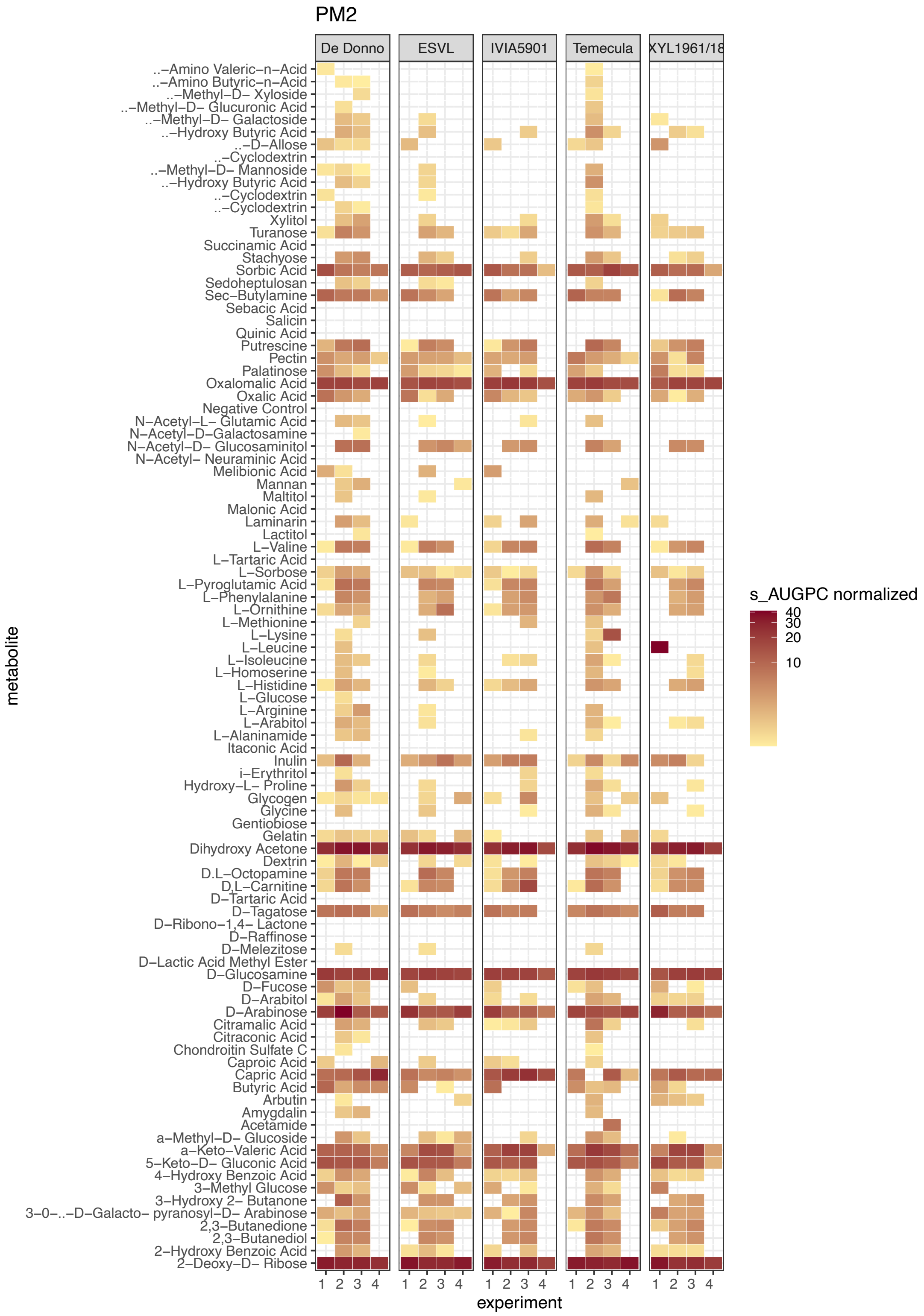


**Figure S6. Bacterial growth in the Biolog Phenotype MicroArray plates PM2 for five *Xylella fastidiosa* strains** (De Donno, ESVL, IVIA5901, Temecula1, and XYL1961/18) used in this study. Four independent experiments were performed for each strain. Growth is presented as the normalized standardized area under the growth curve (sAUGPC). Data is visualized using a pseudo-logarithmic scale.

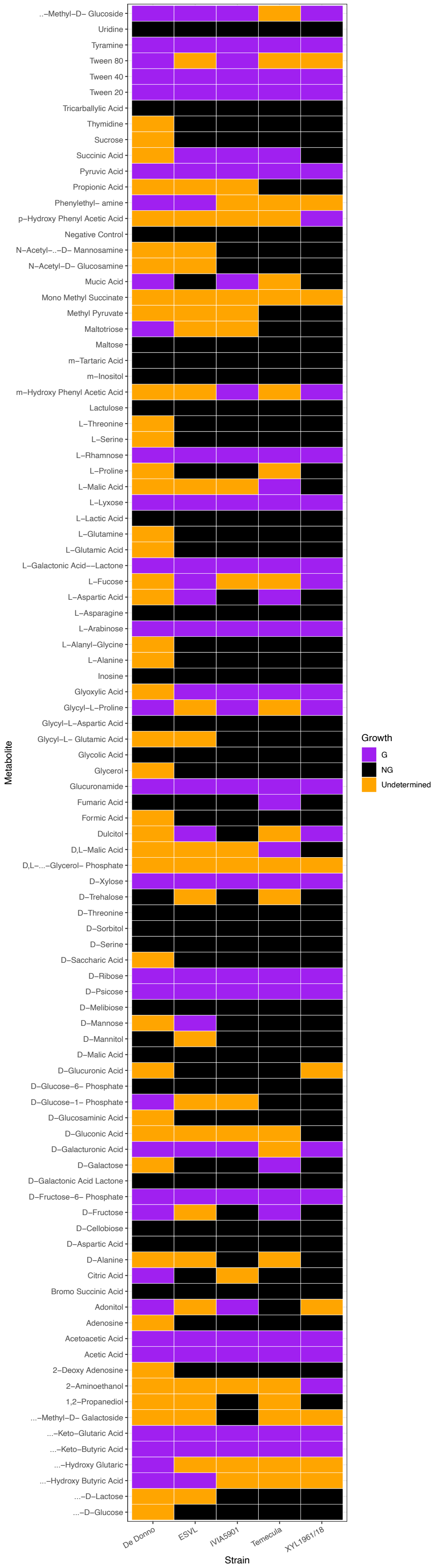

..−Methyl−D− Glucoside
Uridine
Tyramine
Tween 80
Tween 40
Tween 20
Tricarballylic Acid
Thymidine
Sucrose
Succinic Acid
Pyruvic Acid
Propionic Acid
Phenylethyl− amine
p−Hydroxy Phenyl Acetic Acid
Negative Control
N−Acetyl−..−D− Mannosamine
N−Acetyl−D− Glucosamine
Mucic Acid
Mono Methyl Succinate
Methyl Pyruvate
Maltotriose
Maltose
m−Tartaric Acid
m−Inositol
m−Hydroxy Phenyl Acetic Acid
Lactulose
L−Threonine
L−Serine
L−Rhamnose
L−Proline
L−Malic Acid
L−Lyxose
L−Lactic Acid
L−Glutamine
L−Glutamic Acid
L−Galactonic Acid−−Lactone
L−Fucose
L−Aspartic Acid
L−Asparagine
L−Arabinose
L−Alanyl−Glycine
L−Alanine
Inosine
Glyoxylic Acid
Glycyl−L−Proline
Glycyl−L−Aspartic Acid
Glycyl−L− Glutamic Acid
Glycolic Acid
Glycerol
Glucuronamide
Fumaric Acid
Formic Acid
Dulcitol
D,L−Malic Acid
D,L−...−Glycerol− Phosphate
D−Xylose
D−Trehalose
D−Threonine
D−Sorbitol
D−Serine
D−Saccharic Acid
D−Ribose
D−Psicose
D−Melibiose
D−Mannose
D−Mannitol
D−Malic Acid
D−Glucuronic Acid
D−Glucose−6− Phosphate
D−Glucose−1− Phosphate
D−Glucosaminic Acid
D−Gluconic Acid
D−Galacturonic Acid
D−Galactose
D−Galactonic Acid Lactone
D−Fructose−6− Phosphate
D−Fructose
D−Cellobiose
D−Aspartic Acid
D−Alanine
Citric Acid
Bromo Succinic Acid
Adonitol
Adenosine
Acetoacetic Acid
Acetic Acid
2−Deoxy Adenosine
2−Aminoethanol
1,2−Propanediol
...−Methyl−D− Galactoside
...−Keto−Glutaric Acid
...−Keto−Butyric Acid
...−Hydroxy Glutaric
...−Hydroxy Butyric Acid
...−D−Lactose
...−D−Glucose
Metabolite
De Donno
ESVL
IVIA5901
Temecula
XYL1961/18
Strain
Growth
G
NG
Undetermined

**Figure S7. Bacterial growth in the Biolog Phenotype MicroArray plates PM1 for five *Xylella fastidiosa* strains** (De Donno, ESVL, IVIA5901, Temecula1, and XYL1961/18) used in this study. Discrete heatmaps summarize growth categories across the four biological replicates. Growth was categorized based on consistency: a compound was considered assimilated (G) only if the normalized sAUGPC remained positive in at least three of the four biological replicates. Conversely, it was classified as “No growth” (NG) if assimilation occurred in zero or one replicate; otherwise, growth was classified as “Undetermined”.

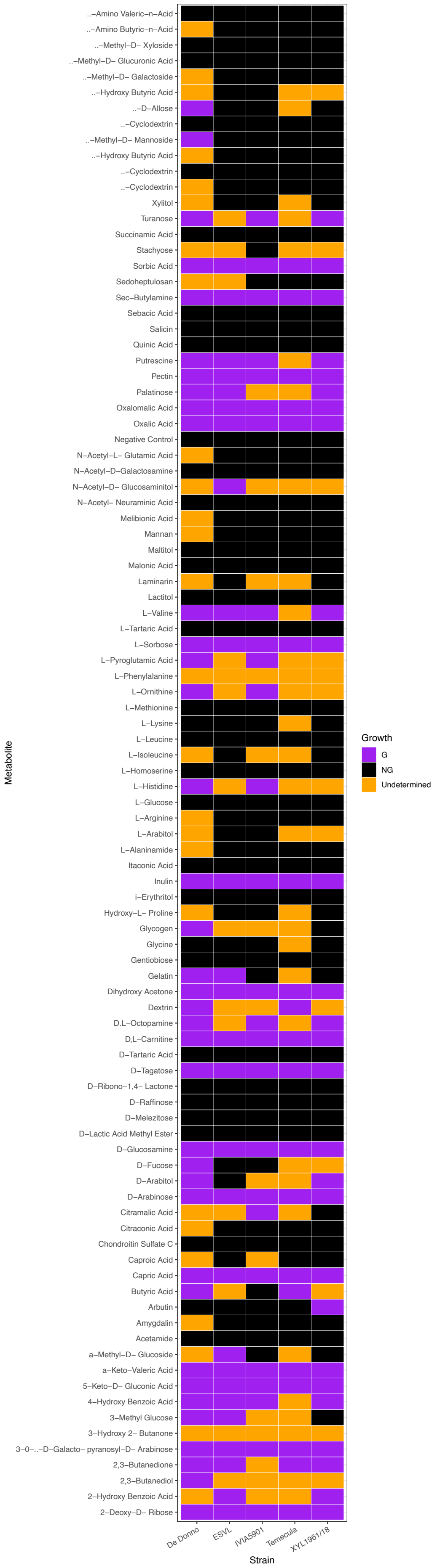
..–Amino Valeric–n–Acid
..–Amino Butyric–n–Acid
..–Methyl–D– Xyloside
..–Methyl–D– Glucuronic Acid
..–Methyl–D– Galactoside
..–Hydroxy Butyric Acid
..–D–Allose
..–Cyclodextrin
..–Methyl–D– Mannoside
..–Hydroxy Butyric Acid
..–Cyclodextrin
..–Cyclodextrin
Xylitol
Turanose
Succinamic Acid
Stachyose
Sorbic Acid
Sedoheptulosan
Sec–Butylamine
Sebacic Acid
Salicin
Quinic Acid
Putrescine
Pectin
Palatinose
Oxalomalic Acid
Oxalic Acid
Negative Control
N–Acetyl–L– Glutamic Acid
N–Acetyl–D–Galactosamine
N–Acetyl–D– Glucosaminitol
N–Acetyl– Neuraminic Acid
Melibionic Acid
Mannan
Maltitol
Malonic Acid
Laminarin
Lactitol
L–Valine
L–Tartaric Acid
L–Sorbose
L–Pyroglutamic Acid
L–Phenylalanine
L–Ornithine
L–Methionine
L–Lysine
L–Leucine
L–Isoleucine
L–Homoserine
L–Histidine
L–Glucose
L–Arginine
L–Arabitol
L–Alaninamide
Itaconic Acid
Inulin
i–Erythritol
Hydroxy–L– Proline
Glycogen
Glycine
Gentiobiose
Gelatin
Dihydroxy Acetone
Dextrin
D.L–Octopamine
D,L–Carnitine
D–Tartaric Acid
D–Tagatose
D–Ribono–1,4– Lactone
D–Raffinose
D–Melezitose
D–Lactic Acid Methyl Ester
D–Glucosamine
D–Fucose
D–Arabitol
D–Arabinose
Citramalic Acid
Citraconic Acid
Chondroitin Sulfate C
Caproic Acid
Capric Acid
Butyric Acid
Arbutin
Amygdalin
Acetamide
a–Methyl–D– Glucoside
a–Keto–Valeric Acid
5–Keto–D– Gluconic Acid
4–Hydroxy Benzoic Acid
3–Methyl Glucose
3–Hydroxy 2– Butanone
3–0–..–D–Galacto– pyranosyl–D– Arabinose
2,3–Butanedione
2,3–Butanediol
2–Hydroxy Benzoic Acid
2–Deoxy–D– Ribose
Metabolite
De Donno
ESVL
IVIA5901
Temecula
XYL1961/18
Strain
Growth
G
NG
Undetermined

**Figure S8. Bacterial growth in the Biolog Phenotype MicroArray plates PM2 for five *Xylella fastidiosa* strains** (De Donno, ESVL, IVIA5901, Temecula1, and XYL1961/18) used in this study. Discrete heatmaps summarize growth categories across the four biological replicates. Growth was categorized based on consistency: a compound was considered assimilated (G) only if the normalized sAUGPC remained positive in at least three of the four biological replicates. Conversely, it was classified as “No growth” (NG) if assimilation occurred in zero or one replicate; otherwise, growth was classified as “Undetermined”.

**Table S1. Composition of the six predicted minimal media (mXC) based on the iXfcore metabolic model**. After predicting a minimal medium that achieved a specific growth rate of 0.413 $h^{-1}$, two initial minimal media (mXC1 and mXC2) were formulated, using different iron sources with glutamine as carbon and nitrogen source. For subsequent media (mXC3 to mXC6), glutamine was replaced with equivalent quantities of alternative carbon sources that supported growth in silico and in vitro, supplementing with specific nitrogen sources and cofactors. All component concentrations were adapted from existing synthetic media known to support *Xylella fastidiosa* growth.

| Component ($g \cdot L^{-1}$) | **mXC1** (glutamine + iron) | **mXC2** (glutamine + hemin) | **mXC3** (acetate + ammonium) | **mXC4** (acetate + ammonium + biotin) | **mXC5** (ribose + ammonium) | **mXC6** (ribose + cysteine) |
|---|---|---|---|---|---|---|
| **Carbon source** | | | | | | |
| L-glutamine | 4 | 4 | - | - | - | - |
| D-Ribose | - | - | - | - | 4.1 | 4.1 |
| Sodium acetate | - | - | 5.6 | 5.6 | - | - |
| **Iron source** | | | | | | |
| Ferric pyrophosphate | 0.25 | - | - | - | - | - |
| Hemin Chloride | - | 0.01 | 0.01 | 0.01 | 0.01 | 0.01 |
| **Inorganic salts** | | | | | | |
| $K_2HPO_4$ | 1.2 | 1.2 | 1.2 | 1.2 | 1.2 | 1.2 |
| $KH_2PO_4$ | 1 | 1 | 1 | 1 | 1 | 1 |
| $MgSO_4$ $7H_2O$ | 0.4 | 0.4 | 0.4 | 0.4 | 0.4 | 0.4 |
| **Nitrogen source** | | | | | | |
| $(NH_4)_2SO_4$ | - | - | 1.32 | 1.32 | 1.32 | - |
| L-Cysteine | - | - | - | - | - | 0.4 |
| **Vitamins/Cofactors** | | | | | | |
| Biotin | - | - | - | 0.0002 | - | - |

**Table S2. Accessory enzymatic activities included in the iXfcore metabolic model** found to be compromised or missing in a small subset of strains, due to pseudogenization. Genes are represented by their corresponding locus tag on the *Xylella fastidiosa* reference strain 9a5c.

| **Locus tag** | **Functional annotation** | **Strain** |
|---|---|---|
| XF_RS00075 | Coproporphyrinogen III oxidase | Mul034 |
| XF_RS00315 | Putative 4-aminobenzoate synthase | M12 |
| XF_RS02735 | Alkaline phosphatase | De Donno, Salento2, J1a12, 3124 |
| XF_RS03390 | Undecaprenyldiphospho-muramoylpentapeptide beta-N-acetylglucosaminyltransferase | XYL1961/18 |
| XF_RS03530 | 2-polyprenyl-6-methoxyphenol hydroxylase | Salento2 |
| XF_RS03665 | Dihydrolipoyllysine-residue acetyltransferase | XYL468, CFBP8418 |
| XF_RS03845 | Magnesium and cobalt transport protein CorA | XYL1961/18 |
| XF_RS04280 | Acetylornithine deacetylase | J1a12 |
| XF_RS04290 | N-acetylglutamyl-phosphate reductase | Pr8x, U24D, J1a12 |
| XF_RS04305 | Gamma-glutamyl-phosphate reductase | M12 |
| XF_RS04390 | Glycerol-3-phosphate 1-O-acyltransferase | XYL468, XYL1961/18, CFBP8418 |
| XF_RS04530 | Glucose-6-phosphate dehydrogenase | Pr8x |
| XF_RS04680 | Carbamoyl phosphate synthase large subunit | Salento2, J1a12 |
| XF_RS04750 | Methylenetetrahydrofolate reductase | J1a12 |
| XF_RS05260 | Fe/S-dependent 2-methylisocitrate dehydratase AcnD | Temecula1, M23, 9a5c, U24D, Ann1 |
| XF_RS05450 | 2-C-methyl-D-erythritol 4-phosphate cytidylyltransferase | XYL468, XYL1961/18, CFBP8418 |
| XF_RS05620 | 3-dehydroquinate synthase | Ann1, U24D, J1a12 |
| XF_RS05675 | Sulfate/thiosulfate import ATP-binding protein CysA | J1a12 |
| XF_RS05835 | Glycine dehydrogenase | Hib4 |
| XF_RS05895 | Phosphoenolpyruvate--protein phosphotransferase | J1a12 |
| XF_RS05995 | Phosphoribosylformylglycinamidine synthase | Pr8x, Salento2, Hib4, J1a12 |
| XF_RS06330 | Sulfite reductase (flavoprotein subunit) | U24D, J1a12 |
| XF_RS06440 | General secretion pathway protein GspK | J1a12 |
| XF_RS06555 | Oxoglutarate dehydrogenase | 3124 |
| XF_RS07865 | 3-oxoacyl-ACP synthase III | J1a12 |
| XF_RS07885 | Acetolactate synthase (large subunit) | Salento2 |
| XF_RS07975 | Glutamine synthetase | SandyiAnn1 |
| XF_RS07985 | Ammonia channel protein | SandyiAnn1 |
| XF_RS08165 | Phosphomethylpyrimidine synthase | XYL1961/18 |
| XF_RS08340 | L-aspartate oxidase | Mul0034 |
| XF_RS08345 | Nicotinate-nucleotide diphosphorylase | Mul0034 |
| XF_RS09080 | Glutamate dehydrogenase | U24D, J1a12 |
| XF_RS09420 | Nicotinate-nicotinamide nucleotide adenylyltransferase | Mul0034 |
| XF_RS09650 | Histidinol-phosphate aminotransferase | Salento2 |
| XF_RS09805 | Acetyl-coenzyme A synthetase | J1a12 |
| XF_RS10000 | Porphobilinogen synthase | IVIA5901 |

| Locus tag | Functional annotation | Strain |
|---|---|---|
| XF_RS10205 | 5'-methylthioadenosine phosphorylase | U24D, 9a5c |
| XF_RS10260 | GumF protein | U24D, Salento2 |
| XF_RS10445 | HMBPP reductase | Salento2 |
| XF_RS10630 | Cytochrome c biogenesis ATP-binding export protein CcmA | XYL1961/18 |
| XF_RS10635 | Heme exporter protein CcmB | Hib4 |
| XF_RS11515 | Ribose-phosphate diphosphokinase | Salento2 |
| XF_RS11745 | Glutamate synthase large subunit | Salento2 |

**Table S3. Enzymatic steps involved in the acetate assimilation pathway predicted by the iXfcore metabolic model.** The pathway enabling acetate into biomass combines two modules: the initial half of the 3-hydroxypropionate bicycle (Herter et al., 2002; steps 1-7), and the methylcitrate pathway for propionyl-CoA assimilation (Brämer & Steinbüchel, 2001; Dolan et al., 2018; steps 8-11). For each enzymatic step represented in Figure 3 (1–11), the table lists the corresponding enzyme, EC number, associated gene(s) identified in *X. fastidiosa*, and the reaction ID in the iXfcore model.

| Step | EC number | Enzyme | Locus tag | Reaction ID |
|---|---|---|---|---|
| 1 | 6.2.1.1 | Acetate-CoA ligase | XF_RS09805 | rxn00175 |
| 2 | 6.4.1.2 | Acetyl-CoA carboxylase (requires biotin) | XF_RS00200 and XF_RS00205 and XF_RS00860 and XF_RS06185 | rxn00533 |
| 3 | 1.2.1.75 | Malonyl-CoA reductase (NADPH) | XF_RS05765 | rxn00532 |
| 4 | 1.1.1.298 | 3-hydroxypropionate dehydrogenase (NADP+) | XF_RS00605 | rxn16149 |
| 5 | 6.2.1.36 | Acyl-CoA synthetase | XF_RS09805 | rxn16146 |
| 6 | 4.2.1.116 | Enoyl-CoA hydratase | XF_RS04720 | rxn02181 |
| 7 | 1.3.1.84 | Acryloyl-CoA reductase (NADPH) | XF_RS07485 or XF_RS10335 | rxn00668 |
| 8 | 2.3.3.5 | Methylcitrate synthase PrpC | XF_RS06495 | rxn00679 |
| 9 | 4.2.1.117 | 2-methylcitrate dehydratase AcnD | XF_RS05260 | rxn03060 |
|  | 5.3.3.7 | 2-methylaconitate cis-trans isomerase PrpF | XF_RS05265 |  |
| 10 | 4.2.1.3 / 4.2.1.99 | Aconitase (AcnB)/2-methylcitrate dehydratase | XF_RS01240 | rxn03061 |
| 11 | 4.1.3.30 | Methylisocitrate lyase (PrpB) | XF_RS05225 | rxn00289 |

**Table S4. Comparison of pathways for acetate assimilation**. The five previously known acetate assimilation strategies as described in the literature (see Schada von Borzyskowski et al. (2020) and Petushkova et al. (2021) for reviews) are the glyoxylate cycle, the ethylmalonyl CoA pathway (EMCP), the methylaspartate cycle (MaC), the ferredoxin-dependent pyruvate synthase (FPS), and half of the 3-hydroxypropionate (3HP) bi-cycle. For the comparison, the stoichiometries of all the pathways were adjusted for the synthesis of 1 oxaloacetate (OAA), including the *X. fastidiosa*'s Xf pathway, albeit the final carboxylation step from pyruvate to OAA is missing in the metabolism of this bacterium. A positive sign denotes the generation of the given compound, while a negative sign means its consumption.

| Pathway | Number of enzymatic steps[a] | Reduced coenzymes produced | Reducing power demand | ATP produced[b] | Energy cost[c] | C co-assimilation |
|---|---|---|---|---|---|---|
| **Glyoxylate cycle** 2 aceCoA = 1 OAA | 2 (+5) | +2 NADH +1 $FADH_2$ | 0 | 6.5 | +2 CoA | 0 |
| **EMCP** 3 aceCoA + $CO_2$ + $HCO_3^-$ = 2 OAA | 14 (+5) | +1 NADH +1 $FADH_2$ | –1 NADPH | 4 | +1.5 CoA | –0.5 $CO_2$ –0.5 $HCO_3^-$ |
| **MaC** 2 aceCoA = 1 OAA | 9 (+4) | +1 NADH +1 $FADH_2$ | –2 NADPH | 4 | +2 CoA | 0 |
| **FPS** aceCoA + $CO_2$ + $HCO_3^-$ = 1 OAA | 1 (+1) | 0 | –1 $Fd_{red}$ | 0 | –1 ATP +1 CoA | –1 $CO_2$ –1 $HCO_3^-$ |
| **Half 3HP bi-cycle** aceCoA + 2 $HCO_3^-$ = 1 OAA | 9 (+4) | +1 NADH +1 $FADH_2$ | –3 NADPH | 4 | -4 ATP +2 CoA | –2 $HCO_3^-$ |
| **Xf pathway** aceCoA + $CO_2$ + $HCO_3^-$ = 1 OAA | 11 (+4) | +1 NADH +1 $FADH_2$ | –3 NADPH | 4 | -4 ATP +2 CoA | –1 $CO_2$ –1 $HCO_3^-$ |

[a] In brackets, additional TCA cycle enzymes.
[b] ATP equivalents assuming aerobic respiration (2.5 ATP/NADH and 1.5 ATP/$FADH_2$)
[c] ATP equivalents and thioester bonds consumed (CoA produced).

**Table S5. Definition of in silico exchange constraints applied for the iXfcore metabolic model under different simulated growth conditions.** Lower and upper bounds are expressed in mmol·gDW$^{-1}$·h$^{-1}$. Specific carbon and nitrogen uptake constraints varied depending on the simulation type (minimal medium prediction, phenotypic substrate evaluation, or trade-off simulations) as described in the *In silico constraints and growth conditions* section.

| Reaction ID | Metabolite | Lower bound (mmol·gDW$^{-1}$·h$^{-1}$) | Upper bound (mmol·gDW$^{-1}$·h$^{-1}$) |
|---|---|---|---|
| **Inorganic ions**[a] | | | |
| EX_cpd00011_e0 | $CO_2$ | -1000 | 1000 |
| EX_cpd00001_e0 | $H_2O$ | -1000 | 1000 |
| EX_cpd00067_e0 | H_plus | -1000 | 1000 |
| EX_cpd00063_e0 | $Ca_2$_plus | -1000 | 1000 |
| EX_cpd00099_e0 | Cl | -1000 | 1000 |
| EX_cpd00149_e0 | $Co_2$ | -1000 | 1000 |
| EX_cpd00058_e0 | $Cu_2$ | -1000 | 1000 |
| EX_cpd10515_e0 | $Fe_2^+$ | -1000 | 1000 |
| EX_cpd10516_e0 | $Fe_3^+$ | -1000 | 1000 |
| EX_cpd00205_e0 | K_plus | -1000 | 1000 |
| EX_cpd00254_e0 | Mg | -1000 | 1000 |
| EX_cpd00030_e0 | $Mn_2$_plus | -1000 | 1000 |
| EX_cpd00971_e0 | Na_plus | -1000 | 1000 |
| EX_cpd00009_e0 | Phosphate | -1000 | 1000 |
| EX_cpd00048_e0 | Sulfate | -1000 | 1000 |
| EX_cpd00034_e0 | $Zn_2$_plus | -1000 | 1000 |
| **Aerobic conditions** | | | |
| EX_cpd00007_e0 | EX_$O_2$_e0 | -1000 | 1000 |
| **Nitrogen source**[b] | | | |
| EX_cpd00013_e0 | $NH_3$ | -1000 (0) | 1000 |
| EX_cpd00023_e0 | L-Glutamate | -6 | 1000 |
| **Carbon source**[c] | | | |
| EX_cpd00053_e0 | L-Glutamine | -7.25 (-15.665) | 1000 |

| Reaction ID | Metabolite | Lower bound (mmol·gDW$^{-1}$·h$^{-1}$) | Upper bound (mmol·gDW$^{-1}$·h$^{-1}$) |
|---|---|---|---|
| x | Substrate | −100 C-mmol·gDW$^{-1}$·h$^{-1}$ | 1000 |

a Inorganic ions were left unconstrained (−1000 to 1000 mmol·gDW$^{-1}$·h$^{-1}$) in all simulations.

b Nitrogen sources were changed depending on simulation: ammonia was set to 0 in the results of the minimal medium and in the trade-off polyamine and virulence factors simulations. Glutamate uptake was set to -6 mmol·gDW$^{-1}$·h$^{-1}$ for phenotypic substrate simulations, including the acetate results.

c Glutamine uptake was constrained to -7.25 mmol·gDW$^{-1}$·h$^{-1}$ for minimal medium prediction, and to -15.665 mmol·gDW$^{-1}$·h$^{-1}$ for trade-offs simulations. Carbon source uptake (x: Reaction IDs in Supplementary Material S8) for phenotypic simulations was normalized and fixed at −100 C-mmol·gDW$^{-1}$·h$^{-1}$.

**Table S6. Percentage of substrates metabolized in Biolog Phenotype MicroArray (PM1 and PM2) plates** by the five *X. fastidiosa* strains used in this study.

| Strain | PM1 | PM2 |
|---|---|---|
| De Donno | 31.6 | 38.9 |
| ESVL | 28.4 | 28.4 |
| IVIA5901 | 27.4 | 27.4 |
| Temecula1 | 26.3 | 21.1 |
| XYL1961/18 | 27.4 | 28.4 |

**Table S7. Comparison between in silico growth predictions and in vitro phenotypic profiles of *X. fastidiosa*.** Experimental growth across various substrates, including carbohydrates, organic acids, lipids, and amino acids, was obtained from Biolog PM plates. In silico growth was predicted through FBA using the iXfcore model as described in Supplementary Material and Methods.

| Substrate | Growth assessment | |
|---|---|---|
| | In silico[a] | Experimental |
| **Carbohydrates and derivatives** | | |
| D-Ribose | + | + |
| D-Xylose | + | + |
| L-Arabinose | - | + |
| L-Lyxose | - | + |
| 2-Deoxy-D-Ribose | - | + |
| D-Arabinose | n.a. | + |
| D-Fructose-6-Phosphate | + | + |
| D-Psicose | - | + |
| D-Tagatose | - | + |
| L-Sorbose | n.a. | + |
| L-Rhamnose | n.a. | + |
| Inulin | n.a. | + |
| Pectin | n.a. | + |
| D-Glucosamine | + | + |
| 3-0-β-D-Galactopyranosyl-D-Arabinose | n.a. | + |
| Glucuronamide | n.a. | + |
| L-Galactonic Acid-γ-Lactone | n.a. | + |
| 5-Keto-D-Gluconic Acid | n.a. | + |
| **Organic acids** | | |
| ∝-Keto-Glutaric Acid | + | + |
| ∝-Keto-Butyric Acid | - | + |
| ∝-Keto-Valeric Acid | n.a. | + |

| Substrate | Growth assessment | |
|---|---|---|
| | In silico[a] | Experimental |
| Pyruvic acid | + | + |
| Acetoacetic Acid | - | + |
| Acetic Acid | + | + |
| Oxalic Acid | n.a. | + |
| Oxalomalic Acid | n.a. | + |
| Dihydroxy Acetone | - | + |
| Sorbic Acid | n.a. | + |
| **Lipids and fatty acids** | | |
| Capric acid | n.a. | + |
| Tween 20 | n.a. | + |
| Tween 40 | n.a. | + |
| D,L-Carnitine | n.a. | + |
| **Amino acids and amines** | | |
| Tyramine | n.a. | + |
| Sec-Butylamine | n.a. | + |

[a] (+) and (-) indicate, respectively, growth/no growth in the corresponding simulation or experimental result. (n.a.) indicates metabolites not represented in the metabolic reconstruction or not available to model.
