## Supplementary Methods for "A metabolic model based on a pangenome core unveils new biochemical features of the phytopathogen *Xylella fastidiosa*"

#### 1. Metabolic model refinement

To facilitate thorough curation of the draft reconstruction, reactions, metabolites, and genes were annotated using identifiers from multiple biochemical databases, including KEGG (Kanehisa et al., 2025), MetaCyc (Caspi et al., 2020), BioCyc (Karp et al., 2019) BiGG Models (King et al., 2016), ModelSEED (Overbeek et al., 2005; Henry et al., 2010), and NCBI databases. SBOannotator (Leonidou et al., 2023) was used to assign SBO terms. BioCyc (Karp et al., 2019), KEGG (Kanehisa et al., 2025), BRENDA (Chang et al., 2021) and Pfam databases (Mistry et al., 2021) were consulted to support manual curation. Consistency analysis of the topological network was conducted to identify gap metabolites and blocked reactions (Ponce-de-Leon et al., 2015). During this step, reaction stoichiometries were revised, including mass and charge balances, reaction directionality and cofactor usage. Similarity-based searches were used to infer missing functions in the model to sequences from the *X. fastidiosa* pangenome, based on sequence and functional homology with other organisms. These analyses were performed using BLASTp (Camacho et al., 2009) and HMMER (Mistry et al., 2013) toolboxes against local sequence databases built from the *X. fastidiosa* pangenome whole-genome sequences. Within the HMMER framework, hmmscan and hmmsearch and pfamscan (Madeira et al., 2024) were used to identify conserved protein domains and domain architectures, providing additional support for functional inference based on profile hidden Markov models. Although all pseudogenes were initially excluded from the pangenome analysis–defined as genes predicted to have lost functionality due to frameshift mutations, partial or incomplete sequences, internal codon stops or a high proportion of ambiguous nucleotides–this criterion was later revised in order to avoid overlooking any essential metabolic activity, during the manual model curation. To that end, every missing function was also searched within the accessory gene sets. Accessory functions that were found in the majority of strains and pseudogenized only in a limited number of strains, typically due to point mutations visualized with the MEGAX software (Kumar et al., 2018), were treated as core functions to better represent the general metabolic phenotype of the species. Reactions were assigned a confidence score ranging from 0 to 4 according to the level of supporting evidence, following the criteria defined by Thiele and Palsson (2010). Curation efforts focused on pathways central to the conserved physiology of *X. fastidiosa*, including central carbon metabolism, complete biosynthetic routes for amino acids, lipid and nucleotide metabolism, energy metabolism, and selected transport systems (Simpson et al., 2000; Meidanis et al., 2002). In addition, pathways related to redox balance, cofactor biosynthesis and host-associated growth were carefully curated. These included quinone and shikimate biosynthesis, polyamine metabolism, the S-adenosylmethionine (SAM) cycle, sulfur and nitrogen metabolism, and the methionine salvage pathway. Vitamin and cofactor biosynthetic pathways were curated in detail, encompassing pantothenate, folate, riboflavin, thiamine, and porphyrin metabolism. Finally, pathways associated with cell envelope biogenesis and host interaction were refined. This included the biosynthesis of exopolysaccharides (EPS) and lipopolysaccharides (LPS), as well as other fatty acid metabolism. Furthermore, the biosynthesis and export reactions of the LesA protein were incorporated to the model, following previous strain-specific metabolic reconstructions (Gerlin et al., 2020; Oliveira et al., 2023).

### 2. In silico contraints and growth conditions

To evaluate phenotypic substrate utilization data in silico, including acetate growth conditions, carbon substrate uptakes were normalized by the number of carbon per molecule, ensuring an equivalent maximum carbon availability across metabolites. A carbon uptake rate of 100 C-mmol·gDW$^{-1}$·h$^{-1}$ was imposed, following the methodology of Gerlin and coworkers (2020). Ammonia uptake was left unconstrained as the nitrogen source. Additionally, L-glutamate uptake was set to -6 mmol $gDW^{-1}$ $h^{-1}$ to evaluate differences in growth between substrates without limitations imposed by ammonium uptake and anaplerotic reactions of the network.

The complete set of exchange flux constraints in each simulation is provided in Table S5.

#### 3. Phenotypic characterization of *Xylella fastidiosa* strains

Phenotypic characterization of five *X. fastidiosa* strains (ESVL, De Donno, IVIA5901, Temecula1, and XYL1961/18) was performed using the Biolog phenotype microarray (PM) system (Biolog, Hayward, California). The potential utilization of 190 carbon sources was determined using the PM1 and PM2 plates. For PM plate inoculation, bacterial cells were scraped from solid culture plates and resuspended in the IF-0a GN/GP base inoculating fluid (Biolog, Hayward, California) and the bacterial suspension was mixed with the corresponding additive solutions for each PM plate and the Dye Mix G, to reach a final optical density at 600 nm ($OD_{600}$) of 0.08 according to the manufacturer's protocol. Subsequently, 100 µL of a cell suspension of each *X. fastidiosa* strain was added to each well of the PM microplates and incubated at 28 °C during 9 days without agitation. Bacterial growth was assessed by measuring $OD_{600}$ using a microplate absorbance reader i-Mark TM spectrophotometer (Bio-Rad, Hercules, CA, USA). Four independent experiments were conducted for each strain and PM substrate-type microplate.

Growth curves of the different *X. fastidiosa* strains growing in each PM were obtained by accumulating $OD_{600}$ values over time from the date of inoculation. Cumulative growth was then calculated by subtracting the initial net absorbance at the beginning of the experiment from the net absorbance on each subsequent day. The standardized area under the growth progress curve (sAUGPC) was calculated using the trapezoidal integration method standardized by duration of the experiment in days (Simko & Piepho, 2012).

In the PM study, metabolic assimilation by *X. fastidiosa* was determined by normalizing the sAUGPC of each compound against the strain- and plate-specific negative control (NC). Following the criteria of Gerlin and coworkers (2020), a compound was considered as positive carbon source ('Growth') if its sAUGPC was at least 1.14 times higher than the NC. Specifically, a normalized sAUGPC was calculated as 100 x [sAUGPC - (1.14 * $sAUGPC_{NC}$)]. Negative values after normalization were set to zero. Growth was categorized based on consistency: a compound was considered assimilated only if the normalized sAUGPC remained positive in at least three of the four biological replicates. Conversely, it was classified as 'No growth' if assimilation occurred in zero or one replicates; otherwise, growth was assessed to be ‘Undetermined’.

To visualize the metabolic profiles across PM plates, two heatmap approaches were used. First, a quantitative heatmap was generated to represent the normalized sAUGPC values, with a pseudo-logarithmic transformation for the color scale. Values of zero (no growth) were set as transparent. Second, discrete heatmaps were obtained to summarize growth categories across the four biological replicates. All visualizations were produced using ggplot2, ggpubr and ggh4x packages in R.
