## Supplementary Results and Discussion for "A metabolic model based on a pangenome core unveils new biochemical features of the phytopathogen *Xylella fastidiosa*"

#### 1. Some metabolic characteristics of the iXfcore metabolic model

Initial curation efforts on the iXfcore model focused on central metabolism and biomass production. Subsequently, complete biosynthetic routes for amino acids, lipids, nucleotides, energy metabolism, and selected transport systems were manually reviewed. Pathways associated with redox balance, cofactor biosynthesis, and host-associated growth were carefully curated, including quinone and shikimate biosynthesis, polyamine metabolism, the S-adenosylmethionine (SAM) cycle, sulfur and nitrogen metabolism, and the methionine salvage pathway. Vitamin and cofactor biosynthetic pathways were curated in detail, encompassing pantothenate, folate, riboflavin, thiamine, and porphyrin metabolism. Finally, pathways associated with cell envelope biogenesis and host interaction were refined, including exopolysaccharide (EPS) and lipopolysaccharide (LPS) biosynthesis, as well as fatty acid metabolism, and the biosynthesis and export of the LesA protein, following previous strain-specific metabolic reconstructions (Gerlin et al., 2020; Oliveira et al., 2023).

During this iterative process, a set of accessory genes were identified as present in the majority of the strains, and considered relevant for modelling the general metabolic behavior of *X. fastidiosa* (Table S2). These genes were not included into the core set of sequences due to pseudogenization events in specific strains. In several cases, this was the result from frameshift mutations caused by single-nucleotide insertions or deletions within repetitive nucleotide regions, which may allow some enzymatic activity or reflect sequencing artefacts (Cooley & Wright, 2024). For instance, the accessory gene XF_RS10205, encoding 5'-methylthioadenosine phosphorylase, is annotated as a pseudogene in strain 9a5c despite being reviewed in UniProt (Q9PAZ2) as a functional enzyme. In total, 43 accessory genes, annotated as pseudogenes, were incorporated into the metabolic reconstruction. Several cases suggest that these annotations may not reflect true gene loss but sequencing mistakes, as some genes appear intact in alternative genome assemblies of the same strain or contain frameshift mutations in homopolymeric regions (Cooley & Wright, 2024). Moreover, some of these genes belong to otherwise complete metabolic pathways, making unlikely the loss of a single intermediate reaction. For these reasons, we carefully evaluated pseudogene annotations during model curation.

The core gene set of *X. fastidiosa* encoded almost all central metabolic functions, although several specific gaps and distinctive features were identified. The Embden-Meyerhof-Parnas (EMP) and the Entner-Doudoroff (ED) glycolytic pathways were fully represented in the iXfcore model. However, one enzyme from the gluconeogenic pathway–namely, fructose-1,6-bisphosphatase (FBPase)–was not identified in any gene set of the pangenome. This absence would suggest limitations in the performance of gluconeogenesis as pointed out in early studies on *X. fastidiosa* metabolism reporting the absence of FBPase (Facincani et al., 2003). Nevertheless, extensive experimental evidence indicates that *X. fastidiosa* is capable of converting pyruvate into sugars (Davis et al., 1980, 1981; Almeida et al., 2004; Gerlin et al., 2020). As previously proposed by Oliveira et al. (2023), FBPase activity (EC 3.1.3.11) can be functionally replaced by a reversible pyrophosphate-dependent phosphofructokinase (PPi-PFK; EC 2.7.1.90), encoded by the core pangenome gene XF_RS01160. This enzyme, homologous to the PPi-PFK described in *Xanthomonas campestris* (UniProt: B0RP51), is reversible and can therefore catalyze both the forward glycolytic reaction and the reverse gluconeogenic conversion. Accordingly, this reaction was implemented in the model to account for both activities.

The oxidative and non-oxidative branches of the pentose phosphate pathway (PPP) were evaluated in the core gene set. Glucose-6-phosphate dehydrogenase (EC

1.1.1.49)–an enzyme shared with the ED pathway–was present in all strains except Pr8x, where it appears as a pseudogene; nevertheless, this activity was retained in the model to reflect its functional relevance. Two additional activities were absent from the core pangenome: 6-phosphogluconate dehydrogenase (PPP oxidative phase) and transaldolase (TALA, PPP non-oxidative phase). Although the absence of the oxidative branch of the pentose phosphate pathway prevents NADPH generation through this route, reducing power may be supplied by the ED pathway and additional reactions such as the malic enzyme. No transhydrogenase activity was identified in the model. Flux redistribution through the reversible reactions of the PPP non-oxidative branch allows synthesis of essential precursors such as phosphoribosyl pyrophosphate (PRPP). In contrast, the absence of TALA is expected to impair growth on pentoses as sole carbon sources, as it disrupts sedoheptulose-7-phosphate (S7P) metabolism. In previously published models (Oliveira et al., 2023), growth on ribose was enabled by incorporating a sedoheptulose-1,7-bisphosphate (SBP) bypass reaction that converts S7P into SBP using PPi instead of ATP, and subsequently cleaved into dihydroxyacetone phosphate (DHAP) and erythrose-4-phosphate (E4P). Experimental and computational studies have shown that bacteria lacking TALA can metabolize S7P through an SBP aldolase pathway (Nakahigashi et al., 2009; Koendjbiharie et al., 2020; Garschagen et al., 2021). For the phosphorylation step, we assigned this activity to the pyrophosphate-dependent phosphofructokinase (PPi-PFK; XF_RS01160), which has been shown in other bacteria to catalyze reversible PPi-dependent phosphoryl transfer reactions (Koendjbiharie et al., 2020). For the aldolase reaction, which converts SBP to DHAP and E4P, we hypothesized that the fructose-bisphosphate aldolase encoded by XF_RS03490 may exhibit substrate promiscuity and catalyze both FBP and SBP cleavage. Such dual specificity has been described in other bacteria, including *Synechocystis* sp. PCC 6803, where class I and II aldolases can participate in SBP metabolism (MetaCyc).

Every enzymatic step of the tricarboxylic acid (TCA) cycle was represented within the core pangenome. The oxoglutarate dehydrogenase complex (ODC) was supported by all its subunits, although the E1 component (XF_RS06555) was absent in strain 3124. The E2 component (XF_RS06550) was conserved in the core genome. The pyruvate dehydrogenase complex (PDC), which catalyzes an analogous reaction and shares a similar multienzyme architecture with ODC, showed strain-specific variation in the E2 subunit. The gene encoding the E2 component (XF_RS03665; dihydrolipoyllysine-residue acetyltransferase) was annotated as a pseudogene in strains XYL468 and CFBP8418, however, inconsistencies between genome assemblies in NCBI suggest potential sequencing or annotation artefacts. So, this complex was conserved in the metabolic model. Two candidate genes encoding the E3 subunit (dihydrolipoyl dehydrogenase) were identified in the pangenome (XF_RS06540 and XF_RS03660). Only XF_RS06540 was part of the core gene set and was therefore assigned as the shared E3 component for both ODC and PDC in the model. Importantly, no enzymes associated with the glyoxylate shunt were identified in any analyzed strain.

Remarkably, pyruvate carboxylase (EC 6.4.1.1), which catalyzes the ATP-dependent conversion of pyruvate to oxaloacetate (OAA), was not identified in the *X. fastidiosa* pangenome and was therefore not included in the model. No alternative reaction directly linking pyruvate to oxaloacetate was detected. The absence of this anaplerotic step limits the replenishment of tricarboxylic acid (TCA) cycle intermediates when sugars or organic acid compounds are supplied as sole carbon sources. Under these conditions, growth requires TCA intermediates to sustain biosynthetic demands. The reconstructed network lacks sufficient anaplerotic reactions to compensate for carbon withdrawal from the cycle, resulting in growth impairment in silico. To evaluate whether growth could be sustained using plant metabolites, we considered incorporating TCA cycle intermediates or their precursors–e.g., glutamine or glutamate–which could be supplied by the host, to simulate bacterial growth.

As previously described, the core metabolic network of *X. fastidiosa* contained complete biosynthetic pathways for all amino acids. Lipid biosynthesis pathways were also identified and included in the reconstruction, whereas no evidence of lipid degradation pathways–e.g., fatty acids $\boldsymbol{\beta}$-oxidation–was detected in the pangenome. Additionally, no auxotrophies were identified for essential vitamins or cofactors.

As for quinone-dependent electron transport, only the ubiquinone biosynthetic pathway was identified in the pangenome, although it appeared incomplete. No candidate gene was detected for two intermediate reactions corresponding to the decarboxylation and hydroxylation of 3-polyprenyl-4-hydroxybenzoate. Nevertheless, the upstream and downstream steps of the pathway were conserved, and no alternative quinone biosynthetic route was identified. Given the apparent continuity of the pathway and the absence of pseudogenization in adjacent enzymatic steps, it is likely that these missing reactions are catalyzed either by highly divergent enzymes not detected by sequence homology, by promiscuous enzymatic activities, or through non-canonical mechanisms that remain to be characterized.

To account for virulence-associated processes, the iXfcore metabolic model incorporated the reactions required for LesA biosynthesis. The gene responsible for LesA production (XF_RS01515 or XF_RS01510) was present in the core gene set, and the ATP-dependent transporter required for its export was also part of the core pangenome, with the exception of one gene involved in the secretion cluster of strain J1A12 (XF_RS06440, *gspK*). However, as described in other cases, the annotation of this gene as a pseudogene was due to a deletion of a G at position 786 within a repetitive G-rich region.

For EPS biosynthesis, the *gum* operon responsible for EPS synthesis and export was found in the core pangenome, with the exception of the *gumF* gene, which encodes an acetyltransferase responsible for the final step of the biosynthetic pathway. This gene was annotated as a pseudogene in strains U24D and Salento 2; however, since the rest of the operon was present and the pseudogene annotation results from a single point mutation that could represent a sequencing error, EPS production was incorporated into the iXfcore model.

### 2. Phenotypic characterization of *Xylella fastidiosa* strains

The phenotypic characterization of strains ESVL, De Donno, IVIA5901, Temecula1 and XYL1961/18 using the phenotype microarray system of Biolog (Supplementary Material S7, Figs. S5 and S6) showed partially overlapping metabolic fingerprints. Among strains, De Donno showed the highest proportion of assimilated carbon sources (31.6% and 38.9% for PM1 and PM2, respectively), whereas Temecula1 showed the lowest (26.3% and 21.1%, respectively) (Table S6). Overall, the five strains consistently metabolized 34 out of the 190 compounds tested (17.9%): 17 compounds in PM1 and 17 compounds in PM2 plates (Table S7; Fig. S7 and S8). Additionally, five compounds were classified as exhibiting “Undetermined” growth across all strains, due to positive growth in only two of the four biological replicates. In contrast, 50 compounds were not assimilated by any of the strains (Supplementary Material S7; Figs. S7 and S8).

The main groups of compounds consistently utilized as carbon sources by all *X. fastidiosa* strains in the PM1 and PM2 plates included i) carbohydrates and their derivatives, ii) organic acids, iii) lipids and fatty acids, and iv) amino acids derivatives and amines. These in vitro results were compared with in silico growth predictions for each compound, when possible (Table S7, Supplementary Material S8).

In silico predictions were consistent with in vitro results for 7 out of the 34 metabolites that supported growth in vitro (D-ribose, D-xylose, D-fructose-6-phosphate, D-glucosamine, α-ketoglutaric acid, pyruvic acid, and acetic acid) (Table S7, Supplementary Material S8. Nineteen out of the 34 metabolites could not be evaluated using the iXfcore model because they were not included in the reconstructed metabolic network or their metabolism could not be represented. These included all lipids and fatty acids, as well as amino acid derivatives and amines. For the remaining eight metabolites, growth was observed in vitro but could not be supported in silico (L-arabinose, L-lyxose, 2-deoxy-D-ribose, D-psicose, D-tagatose, α-ketobutyric acid, acetoacetic acid, and dihydroxyacetone).

Additionally, several compounds that did not support growth for all strains in the PM plates were evaluated in silico due to their reported abundance in the xylem sap of olive, almond, and citrus trees (Table S7, Supplementary Material S8) (Youssefi et al., 2000; Purcino et al., 2007; Anguita-Maeso et al., 2021; Fausto et al., 2021; Surano et al., 2024). D-glucose, one of the major sugars detected in olive xylem sap, was predicted to sustain growth in silico, consistent with previous strain-specific models (Gerlin et al., 2020; Oliveira et al., 2023). However, this prediction was not supported experimentally, as only one strain showed undetermined growth while the remaining four exhibited no growth in the PM assays (Fig. S7). Citric acid, also reported as a common organic acid in xylem sap, was predicted to sustain growth in silico. The corresponding transporter was present in the core pangenome, and growth has been supported in other reconstructions. Experimentally, citrate supported growth in one strain, showed indeterminate growth in one strain, and resulted in no growth in the remaining three strains.

Regarding amino acids commonly reported in xylem sap (glutamine, glutamate, aspartate, arginine, asparagine, alanine, serine, and proline), in silico simulations predicted growth on L-glutamine, L-glutamate, L-aspartate, and L-alanine (Table S7, Supplementary Material S8). However, experimental validation showed partial or limited agreement (Fig. S7 and S8). L-aspartate supported growth in vitro in two strains, while one strain showed undetermined growth and two showed no growth. In contrast, L-glutamate, L-glutamine, and L-alanine predominantly resulted in undetermined or no-growth phenotypes across strains despite being predicted to sustain growth in silico. L-arginine, L-asparagine, L-serine, and L-proline were not predicted to support growth in silico. Consistently, these amino acids did not sustain robust growth in vitro, with most strains showing either no growth or undetermined phenotypes.

The phenotypic characterization performed in this work provides a robust dataset, including four independent experiments and five strains belonging to the three *X. fastidiosa* subspecies (*fastidiosa*, *multiplex* and *pauca*). Previous studies have reported Biolog results for *X. fastidiosa*, such as the work of Gerlin et al. (2020), although in that case the analysis was limited to the strain CFBP8418 (subsp. *multiplex*) used for metabolic modelling.

Our results revealed some discrepancies between in vitro phenotyping, published observations, and model predictions. Consistent growth on glutamine or glutamate was not detected in the Biolog PM assays for the strains analyzed, whereas growth was sustained in silico. However, glutamine was selected as the main carbon source for the minimal media experiments and simulations because its utilization has been widely reported in the literature for *X. fastidiosa* (Almeida et al., 2004; de Macedo Lemos et al., 2003; Gerlin et al., 2020). In addition, in the minimal media experiments performed in this work, glutamine sustained both bacterial growth and biofilm formation. As per the same reason, glutamic acid was used in the phenotypic simulations because it provided a direct entry point into the TCA cycle, representing a known nitrogen source present in xylem sap (Surano et al., 2024). This glutamate supplementation was consistent with the medium composition used in the Biolog phenotypic assays (available via manufacturer). Moreover, its assimilation is consistent with the predictions of the iXfcore model, which requires substrates capable of replenishing TCA cycle intermediates.

Some carbohydrates and derivatives, as well as some organic acids, supported growth in vitro but not in silico, suggesting that certain metabolites and metabolic reactions may be missing from the model, possibly because the corresponding genes are not present in all strains of the pangenome. In the case of lipid substrates, in silico validation could not be performed because the β-oxidation pathway has not been described in *X. fastidiosa*. However, growth on lipid derivatives has been previously reported (Gerlin et al., 2020), suggesting that some alternative, yet undescribed, metabolic pathways may exist. As shown for acetate assimilation, the metabolic model may help guide the search for such alternative pathways by identifying potential metabolic gaps.

Finally, several compounds commonly present in xylem sap were also evaluated. Although no growth was detected in vitro for some of them (D-glucose, citrate, L-aspartate and L-alanine), the metabolic model predicted that these substrates could theoretically support growth. These discrepancies may be explained by differences in experimental conditions. In this work, growth was evaluated using $OD_{600}$ measurements, whereas Biolog PM assays typically rely on respiration-based detection. Additionally, factors such as substrate concentration, medium composition, or transport limitations may influence the ability of cells to utilize certain compounds in vitro. For instance, a low initial cell density could limit the activity of the glutamate transporter, preventing the flux required to sustain growth, unlike other carbon sources such as pyruvate. Moreover, in xylem sap *X. fastidiosa* is likely exposed to combinations of carbon sources, which could compensate for metabolic limitations that are not captured when substrates are tested individually in Biolog PM assays.

#### 3. A proposed pathway for acetate assimilation in *Xylella fastidiosa*

Here, we describe the predicted metabolic solution for growth on acetate as the sole carbon source in more detail, indicating the specific steps shown in Fig. 3.

First, acetate enters the cell through a sodium-dependent acetate symporter (XF_RS09795), which is present in all strains analyzed. Subsequently, acetate is converted into acetyl-CoA by acetate-CoA ligase (Fig. 3, step 1; XF_RS09805). This gene is absent in the J1A12 strain due to a single nucleotide deletion (loss of an adenine at position 1120) within a repetitive region of adenine. Apart from this frameshift mutation, the sequence is identical to that found in the remaining strains. In step 2, acetyl-CoA carboxylase catalyzes the ATP-dependent carboxylation of acetyl-CoA to produce malonyl-CoA. This enzyme requires biotin as a cofactor for its catalytic activity. Malonyl-CoA is then reduced to malonate semialdehyde (3-oxopropanoate; step 3) by a NADP-dependent malonyl-CoA reductase. Although the associated gene (XF_RS05765) is annotated as aspartate semialdehyde dehydrogenase, BLAST analysis and Pfam domain architecture match the reviewed UniProt entry A4YEN2 from *Metallosphaera sedula*, which performs this specific reaction. Step 4 involves the conversion of malonate semialdehyde to 3-hydroxypropionate, catalyzed by 3-hydroxypropionate dehydrogenase. This reaction was incorporated into the model and assigned to gene XF_RS00605, annotated as NADP-dependent 3-hydroxy acid dehydrogenase. In step 5, 3-hydroxypropionate is converted into its corresponding CoA-activated form by an acyl-CoA synthetase, again encoded by XF_RS09805. In this case, BLAST analysis and Pfam domain architecture matched the reviewed UniProt entry A4YGR1 from *M. sedula*, which catalyzes this specific reaction. This enzyme, annotated as "acs" in several strains, would exhibit substrate flexibility, being also responsible for acetate activation. Step 6 involves the dehydration of 3-hydroxypropionyl-CoA to acryloyl-CoA, catalyzed by an enoyl-CoA hydratase, operating in the reverse direction. Although the candidate gene XF_RS04720 identified did not yield a strong BLAST match to experimentally characterized acryloyl-CoA hydratases, it is annotated as an enoyl-CoA hydratase, consistent with the proposed reaction. Furthermore, Pfam domain analysis revealed the presence of the conserved ECH_1 domain, which is characteristic of enoyl-CoA hydratase family enzymes and is found in all reference proteins associated with this activity. Step 7 corresponds to the final reaction of the 3HP bi-cycle module, involving the conversion of acryloyl-CoA into propionyl-CoA. This reaction is catalyzed by an NADPH-dependent acryloyl-CoA reductase. Two candidate genes belonging to the same orthologous group were identified as potentially responsible for this activity (XF_RS07485 and XF_RS10335), both annotated as NAD(P)-dependent alcohol dehydrogenases. BLAST analysis showed that XF_RS07485 matched the reviewed UniProt entry A4YGN2 from *M. sedula*.

Propionyl-CoA is the metabolic junction between the module of part of 3HP bi-cycle and the putative second module of the acetate assimilation, namely, the methylcitrate cycle. Step 8 corresponds to the condensation of propionyl-CoA with oxaloacetate to form 2-methylcitrate. This reaction is classically catalyzed by methylcitrate synthase (PrpC). No canonical *prpC* gene was identified in the core genome. However, the gene XF_RS06495, which is conserved across all strains, annotated as citrate synthase (GltA), was identified as a potential candidate for this step. Given this structural homology and the absence of a dedicated *prpC* gene, GltA was considered the most parsimonious candidate to catalyze the condensation of propionyl-CoA with oxaloacetate in the reconstructed pathway. Such substrate flexibility has been reported for related enzymes within this family (for instance, *Bacillus subtilis*, UniProt entry P45858, as reported in BRENDA database). Step 9 corresponds to the conversion of 2-methylcitrate into 2-methylaconitate, which proceeds through two consecutive activities: 2-methylcitrate dehydratase (EC 4.2.1.117; AcnD) followed by 2-methylaconitate *cis-trans* isomerase (EC 5.3.3.7; PrpF). These transformations were included in the model as a combined step. The *acnD* locus (XF_RS05260) was annotated as a pseudogene in a subset of genomes (Temecula1, M23, 9a5c, U24D and Ann1). Among the strains used experimentally, only Temecula1 carried this pseudogene annotation, whereas the remaining experimental strains did not show this disruption. How this strain solves the absence of a functional AcnD remains unsolved, and it would be worth studying the substrate promiscuity of other aconitases. The gene *prpF* was identified in the core gene set of the pangenome (XF_RS05265). Step 10 corresponds to aconitase AcnB, present in all strains and encoded by XF_RS01240, and annotated as an aconitate hydratase 2 / 2-methylisocitrate dehydratase. Finally, step 11 corresponds to methylisocitrate lyase (PrpB), which belongs to the core gene set and is encoded by XF_RS05225, which produces pyruvate and succinate.
